# The cosmopolitan appendicularian *Oikopleura dioica* reveals hidden genetic diversity around the globe

**DOI:** 10.1101/2022.08.09.503427

**Authors:** Aki Masunaga, Michael J Mansfield, Yongkai Tan, Andrew W Liu, Aleksandra Bliznina, Paolo Barzaghi, Tamara L Hodgetts, Alfonso Ferrández-Roldán, Cristian Cañestro, Takeshi A Onuma, Charles Plessy, Nicholas M Luscombe

## Abstract

Appendicularian tunicates are some of the most abundant mesozooplankton organisms with key roles in marine trophic webs and global carbon flux. Like most appendicularians with cosmopolitan distributions, *Oikopleura dioica* Fol, 1872 is considered a single species worldwide based on morphological features that distinguish them from other appendicularians. Despite their abundance however, there are still only ∼70 described appendicularian species, compared with over 2,800 ascidian tunicates. Here we perform a molecular phylogenetic, morphological, and reproductive assessment of *O. dioica* specimens collected from the Ryukyu Archipelago, mainland Japan, and Europe. The specimens are morphologically very similar, with only detailed examination of the oikoplastic epithelium and quantitative measurements revealing minor distinguishing characteristics. Phylogenetic analyses of the ribosomal gene loci and mitochondrial cytochrome oxidase I (COI) gene strongly indicate that they form three separate genetic clades despite their morphological similarities. Finally, *in vitro* crosses between the Ryukyu and mainland Japanese specimens show total prezygotic reproductive isolation. Our results reveal that the current taxonomic *O. dioic*a classification likely hides multiple cryptic species, highlighting the genetic diversity and complexity of their population structures. Cryptic organisms are often hidden under a single species name because their morphological similarities make them difficult to disinguish and their correct identification is fundamental to understanding Earth’s biodiversity. *O. dioica* is an attractive model to understand how morphological conservation can be maintained despite pronounced genetic divergence.

## Introduction

Appendicularians or larvaceans (Tunicata: Appendicularia) are abundant holoplanktonic tunicates with a global distribution. They are one of the most abundant taxonomic groups in the zooplankton community (Alldredge 1976; Clarke and Roff 1990; Hopcroft and Roff 1995). Under optimal conditions, appendicularians form rapid blooms (Essenberg 1922; Tokioka 1955) exceeding 53,000 individuals per cubic meter (Uye and Ichino 1995). The animals secrete an intricate structure known as a house, which they use to filter-feed suspended dissolved organic particles and microorganisms from the water (Sato et al. 2001); this allows them to feed on pico- and nanophytoplankton that are often too small to be captured by most zooplankton (Alldredge 1976; Saito 2019). Therefore, they serve as an important intermediate link in the food chain connecting pico- and nanophytoplankton and larger zooplankton including fish larvae (Sakaguchi et al. 2017) copepods (Ohtsuka and Onbé 1989), ctenophores (Shiganova 2005; Purcell et al. 2010), jellyfish, and chaetognaths (Alldredge 1976). Most appendicularians measure from 2 to 8 mm in length, with houses ranging from 4 to 38 mm in diameter (Alldredge 1977). However, *Bathochordaeus*, known as giant larvaceans, have body lengths of ∼10cm and construct houses that exceed 1 m in diameter (Hamner and Robison 1992; Katija et al. 2017). Appendicularians maintain efficient feeding by continuously replacing houses: for instance, a single *Oikopleura dioica* can build approximately 50 houses during its lifecycle (Sato et al. 2001). Discarded houses together with trapped materials form carbon-rich aggregates and are one of the main components of the marine snow that eventually sinks to the seabed. Therefore, appendicularians make a major contribution to vertical global carbon flux by transporting a substantial amount of surface productivity to the deep benthic fauna (Alldredge et al. 2005; Robison et al. 2005).

Despite their abundance and ecological importance in functioning ecosystems, the recorded appendicularian diversity remains low with only ∼70 described species worldwide (Tokioka 1960; Hopcroft 2005; Gorsky and Castellani 2017). This trend is especially apparent when compared with sessile tunicates, ascidians, which comprise over 2,800 described species (Shenkar and Swalla 2011). The comparative lack of knowledge is partly due to the challenge of studying appendicularians. Soft-bodied appendicularians are extremely fragile; they are easily damaged during conventional plankton sampling, leaving little material for morphological identifications (Tokioka 1960). In addition, their distributions are often seasonal (Uye and Ichino 1995; Tomita et al. 2003); therefore, continuous and extensive sampling effort is needed to appreciate their true diversity. Furthermore, there might be potential cryptic speciation among currently recognized species (Hopcroft and Robison 1999; Hopcroft 2005): cryptic species are defined as “two or more distinct species that are erroneously classified (and hidden) under one species name” because they are superficially morphologically indistinguishable (Bickford et al. 2007). They are common in many marine taxa including foraminifera (de Vargas et al. 1999), cnidarians (Dawson and Jacobs 2001), fishes (Borsa 2002), elasmobranchs (Quattro et al. 2006), and copepods, with numerous sibling species retaining strong morphological conservation (Chen and Hare 2011; Blanco-Bercial et al. 2014).

There have been three significant periods of appendicularian taxonomic discovery: exploratory studies led by Lohmann, Bückmann, and Hentschel in the late 19th to early 20th centuries; Japanese exploration by Tokioka in the 1950s; and mesopelagic explorations led by Fenaux, Youngbluth, Hopcroft, Robison, and Flood in the late 20th century. These efforts contributed to the current understanding of appendicularian diversity and distributions (Hopcroft 2005). Most appendicularian species were previously delineated using classical morphology-based taxonomy. However, to assess accurately the true diversity of appendicularians, it is also important to examine molecular data that might uncover genetically distinct lineages hidden behind morphologically conserved populations.

Recent combined morphological and molecular taxonomic analyses have described new species of appendicularians and closely related taxa (Garić and Batistić 2016; Sherlock et al. 2017; Sanchez et al. 2019). The molecular approach provides additional evidence for species identification and enables a more accurate assessment of morphologically conserved taxa. Markers such as the cytochrome c oxidase subunit I (COI) and ribosomal DNA (rDNA) genes are often used to elucidate phylogenetic relationships (Zagoskin et al. 2014). In most eukaryotes including appendicularians, rDNA genes are organized in tandemly repeated clusters, each containing the 18S, 5.8S, and 28S ribosomal genes separated by internal transcribed spacers (ITS1 and ITS2) and an intergenic spacer (Singer and Berg 1991). Phylogenetic relationships at higher systematic levels (orders, families, and genera) are often resolved using the highly conserved 18S and 28S rDNA regions. The less conserved internal spacer sequences enable greater phylogenetic resolution of lower taxonomic groups, including species (Schlötterer et al. 1994; Hwang and Kim 1999; Dyomin et al. 2017).

*Oikopleura dioica* is an appendicularian model that contributes to a wide range of biological studies including developmental biology, evolution, genomics, and ecology (Nishida 2008; Lombard et al. 2009; Troedsson et al. 2013; Deng et al. 2018; Onuma and Nishida 2021; Ferrández-Roldán et al. 2021). Much of our knowledge of appendicularian biology comes from multigenerational culturing of *O. dioica* in the laboratory (Paffenhöfer 1973; Bouquet et al. 2009; Martí-Solans et al. 2015; Masunaga et al. 2020). The organism is mainly characterized by its ubiquitous distribution, dioecious nature, and presence of two subchordal cells on its tail. *O. dioica* is currently considered a single species worldwide; however, there are substantial nucleotide sequence variations between *O. dioica* collected from Norway, Osaka, and Okinawa (Denoeud et al. 2010; Wang et al. 2015a, 2020; Bliznina et al. 2021) despite the low level of phenotypic disparity.

Here we perform a molecular phylogenetic, morphological, and reproductive assessment of *O. dioica* collected from the Ryukyu Archipelago, mainland Japan, and Europe. Our results reveal three genetically distinct lineages despite near-indistinguishable morphologies. We provide compelling evidence that *O. dioica* collected from the Ryukyu Archipelago is a separate species from those found in mainland Japan and Europe.

## Materials and methods

### Origins of O. dioica data

*O. dioica* specimens were either obtained from field collections (Kume, Amami, Nagasaki, Hyogo, Aomori) or laboratory cultures (Okinawa, Osaka, Barcelona). Sequence information for molecular phylogenetic analyses was obtained from specimens, or where publicly available, from GenBank or the supplementary material of the corresponding paper (Norway, Croatia). Images, size measurements, and fertilization rates were obtained using laboratory specimens. Sequencing data for all other appendicularians were obtained from GenBank. Collection sites are shown in Figure 1, and detailed sample information is provided in Supplementary Table S1.

**Fig. 1.**
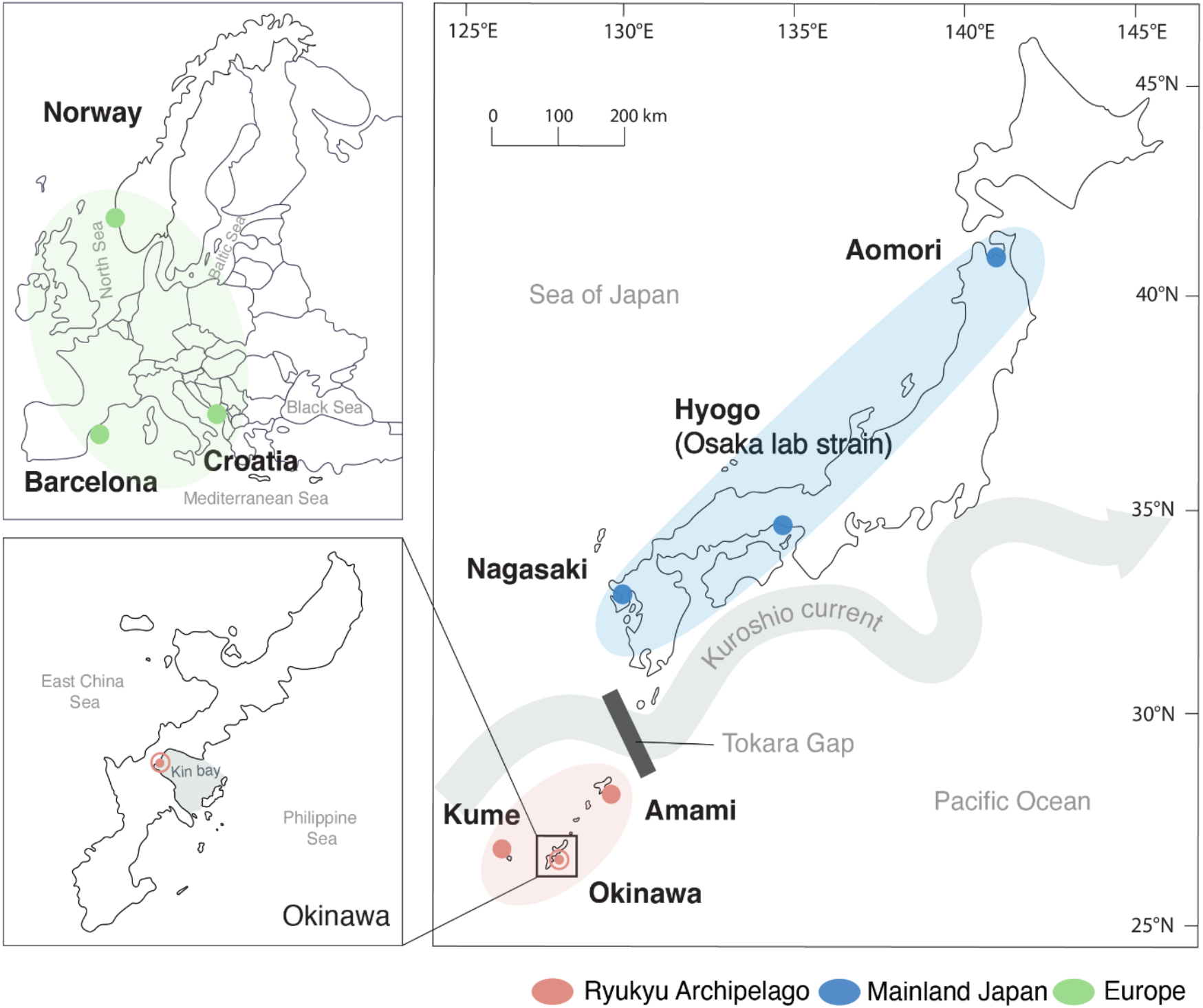
Collection sites of *Oikopleura dioica* used in this study. The Okinawa, Osaka, and Barcelona laboratory strains were initially isolated from the coastlines of Okinawa, Hyogo, and Barcelona

### Field collections of O. dioica from the Ryukyu Archipelago

We surveyed *O. dioica* along the Ryukyu Archipelago from harbors and fishing ports using a hand-held plankton net (100-µm mesh) with a 500 mL volume cod-end (Masunaga et al. 2020). Animals were obtained in June and July 2019 from Kin bay on Okinawa island (26°25’39.6”N 127°49’56.5”E), Kume island (26°21’03.2”N 126°49’17.4”E), and Amami island (28°24’53.6”N 129°35’27.9”E; Fig. 1). *O. dioica* were identified by the presence of two subchordal cells at the distal half of their tails under a dissecting microscope (Fredriksson and Olsson 1991; Fig. 2B). Live animals from Okinawa Island were transported to the Okinawa Institute of Science and Technology Graduate University (OIST) to establish laboratory cultures (Masunaga et al. 2020). Animals from Kume and Amami Islands were isolated and maintained in beakers with portable paddles and motors until they reached maturity on-site. Once the animals matured, males were rinsed three times for 10 min with autoclaved filtered seawater, preserved in RNA*later* (ThermoFisher AM7021), and transported back to OIST.

**Fig. 2.**
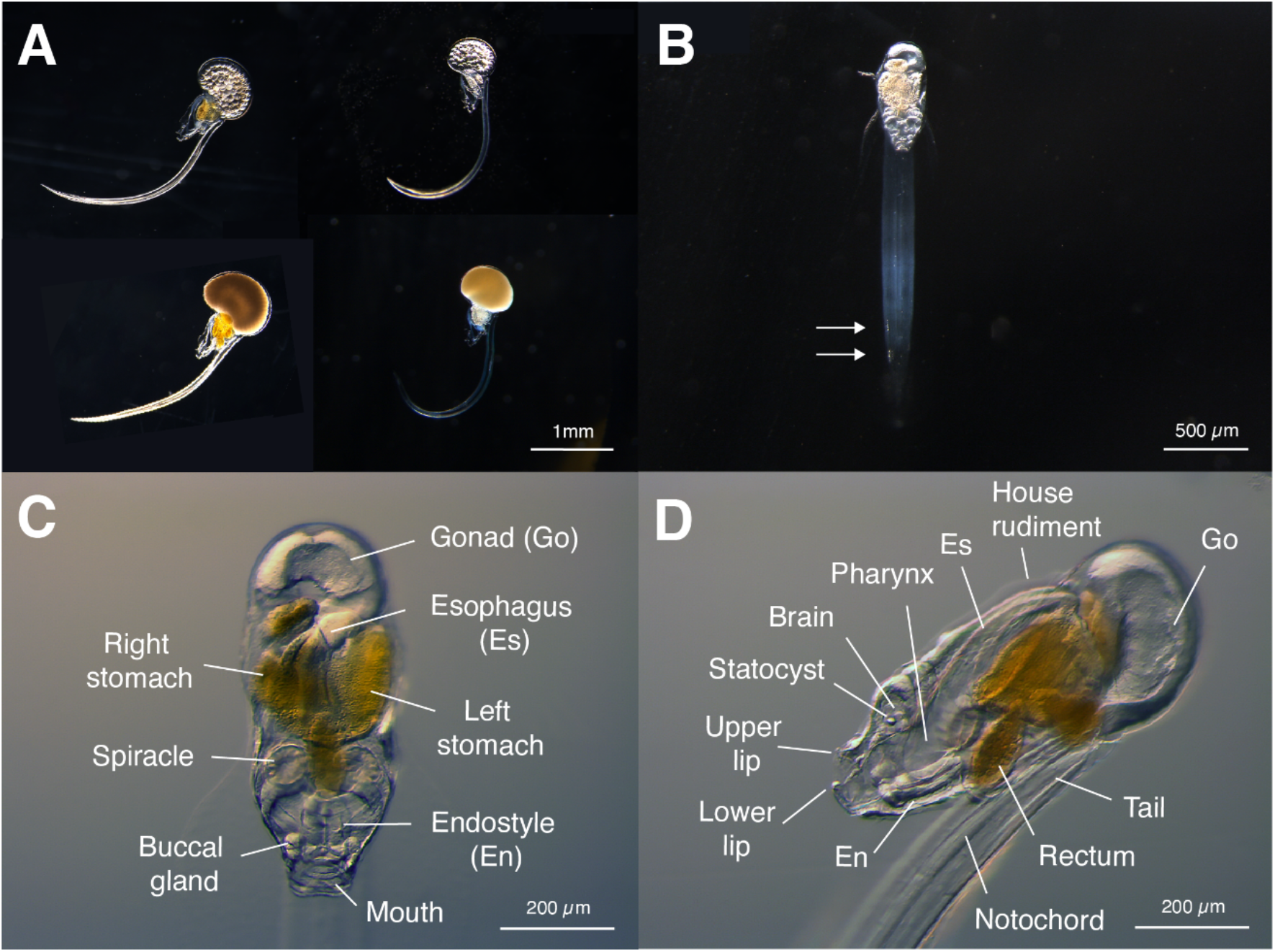
Okinawa and Osaka *O. dioica* laboratory strains. (A) Fully matured females (top) and males (bottom) of Osaka (left column) and Okinawan (right column) *O. dioica* maintained at the OIST culturing facility. (B) Okinawan *O. dioica* individual with two subchordal cells (white arrows) on its tail. (C) Dorsal and (D) ventral views of Okinawan *O. dioica*

### Collections of O. dioica from mainland Japan and Europe

*O. dioica* specimens were generously donated by the Kujukushima Aquarium (UMI KIRARA; Nagasaki), the Nishida lab from Osaka University (Hyogo), the Nishino lab from Hirosaki University (Aomori), and the Cañestro lab from the University of Barcelona (Barcelona). One of the Osaka samples and the Barcelona sample are laboratory cultures of locally collected specimens.

### Laboratory culturing of O. dioica specimens

Live *O. dioica* laboratory animals originally collected from Sakoshi bay in Hyogo prefecture (Osaka lab strain) were kindly provided by the Nishida lab and cultured at the OIST culturing facility. Both Okinawa and Osaka lab strains were cultured at 23°C at OIST (Masunaga et al. 2020). The Barcelona laboratory strain is maintained by the Cañestro lab at the University of Barcelona (Martí-Solans et al. 2015). All animals are fed once or twice a day on a mixture of microalgae and cyanobacteria as previously described (Martí-Solans et al. 2015; Masunaga et al. 2020).

### Genomic DNA isolation and sequencing

Fully mature males were rinsed and either processed immediately for DNA extraction or preserved in RNA*later* for transport. Genomic DNA (gDNA) was isolated using a modified salting-out protocol. First, tissues were lysed with 10 µL/mg proteinase K for 30 min at 56°C. Next, proteins were precipitated out with 50 µL 5M NaCl and separated by centrifugation at 5,000 ×*g* for 15 min at 4°C. The aqueous phase was slowly transferred with a wide-bore pipette to a tube containing 400 µL of cold 100% ethanol and 10 µg/mL glycogen. gDNA was precipitated at −80°C for 20 min and centrifuged at 6,520 ×*g* for 5 min at 4°C. Finally, the pellet was washed with 200 µL of 70% ethanol, allowed to air dry for 5 min, and resuspended in molecular biology grade H2O. gDNA was quantified using a Qubit 3 Fluorometer (Thermo Fisher Scientific, Q32850), and its integrity was measured using Agilent 4200 TapeStation (Agilent, 5067-5365). Isolated gDNA were processed with the Ligation Sequencing Kit (Nanopore LSK109) according to the manufacturer’s protocol and sequenced on the Nanopore MinION platform.

### 18S and ITS phylogenetic analysis

We combined appendicularian 18S ribosomal DNA (rDNA) gene sequences from GenBank with 18S RNA gene sequences identified from our own genome assemblies. All sequences have been deposited in the GenBank database (accession numbers provided in Supplementary Table S1). To identify 18S rDNA genes, we searched genome sequences for the best-hit matches to the Rfam models for the eukaryotic small subunit (SSU; RF01960) and large subunit (LSU; RF02543) using cmsearch of the Infernal package (v.1.1.4). The SSU and LSU models from Rfam were downloaded in April 2021. The internal transcribed spacer (ITS) regions were obtained by extracting the region between SSU and LSU. Multiple sequence alignments for the 18S and ITS regions were created with MUSCLE (v5) within Seaview (v.3.2). Phylogenetic trees were estimated using maximum likelihood with IQ-TREE software (Trifinopoulos et al. 2016) and Bayesian inference with MrBayes (v.3.2.0) (Ronquist and Huelsenbeck 2003). The ML method was calculated using the best model estimated by ModelFinder (GTR+F+I+G4) (Kalyaanamoorthy et al. 2017) according to the Bayesian Information criterion with 1,000 ultrafast bootstrap replicates (Hoang et al. 2018). For Bayesian inference, a GTR substitution model with 6 gamma-distributed rate categories and an uninformative Dirichlet prior was used. Three independent MCMC chains were run with a maximum of 1,000,000 generations and 25% burn-in fraction, allowing the chain to terminate when the standard deviation of split frequencies was lower than 0.01. The 50% majority rule consensus trees with their posterior probabilities are presented.

### COI phylogeny

The mitochondrial cytochrome oxidase I (COI) sequences representing the European lineage were obtained from (Pichon et al. 2019) (PMID: 32148763). The COI sequence representing the mainland Japanese lineage is contig comp28493_c0_seq1_38 from the transcriptome assembly of Wang et al. 2015a (GEO ID: GSE64421, PMID: 26032664). The COI sequence representing the Ryukyu lineage is the transcript model TRINITY_DN9989_c0_g1_i1 from the transcriptome assembly from Bliznina et al. 2021.

### Imaging the oikoplastic epithelium

#### Okinawa and Osaka laboratory strains

To examine the oikoplastic epithelium (OE), immature adults were fixed with 4% paraformaldehyde in 0.1M MOPS and 0.5M NaCl buffer (PFA). Fixed animals were stored at 4°C until use. The animals were washed three times with PBST (PBS with 0.1% Triton X), and nuclei were stained with DAPI. After 2 min incubation at room temperature, the animals were washed three times with PBST and mounted in 30% glycerol. The nuclei of the OE were imaged using a Nikon A1R confocal microscope with a 20x objective. A volume of 40 planes (0.7 micron steps) was observed and two images were averaged for every plane. The volumes were 3D rendered using the NIS Elements imaging software (v.5.21.02) with a 3D LUT (Depth coded Alpha, Altitude). Maximum intensity projections (MIPs) were extracted as a single volume snapshot. Images were post-processed using Adobe Photoshop CC (22.4.1). The number of nuclei in the Fol and the Eisen domains of all specimens were manually identified and counted using the nomenclature described by Kishi et al. (2017).

### Barcelona laboratory strain

Barcelona animals were fixed with 4% paraformaldehyde in 0.1 M MOPS, 0.5 M NaCl, 2 mM MgSO4, 1 mM EGTA. The animals were washed with PBS with 0.1% Tween-20, and nuclei were stained with Hoechst. The animals were mounted in 80% glycerol. The nuclei were imaged using a Zeiss LSM880 microscope with a 25X objective.

### Measuring trunk and tail ratios

Trunk and tail measurements were taken for approximately 50 animals of various developmental stages post-hatching. Each animal was transferred to a Petri dish containing 0.5% agarose. Excess seawater was removed until animals lay flat on the agarose bed and were immobilized. Each animal was photographed (Leica MC 190 HD or BFLY-U3-23S6C) and the trunk length (distal end of gonad to the mouth) and the tail length were measured using the Leica Las software (v.4.9) or FIJI (Schindelin et al. 2012). The trunk-to-tail ratio was calculated as trunk length (mm) / tail length (mm).

### Measuring egg diameters

To compare egg diameters, approximately 20 females were collected from each laboratory strain. Females were rinsed three times with 5 mL of autoclaved filtered seawater (AFSW) to avoid the potential presence of sperm from the cultures. Each female was transferred to a Petri dish and allowed to spawn naturally. Photographs of eggs were taken, upon spawning, and five representative eggs from each clutch were selected for measurements. For each female, we report the mean diameter of five eggs. One-way analysis of variance (ANOVA) was performed on both trunk and tail and egg diameter measurements using R (v.3.6.3, R Core Team 2020).

### Crossing experiment

60 fully mature Okinawa and Osaka animals were randomly assigned to four mating groups during the experimental period of April to May 2018: 10 pairs of Okinawa males and Osaka females, 10 pairs of Okinawa females and Osaka males, and 5 pairs from each strain for positive controls (Okinawa males and females, Osaka males and females). Animals were rinsed three times with 5 mL of AFSW. Each male was transferred to an Eppendorf tube containing 50 µL of AFSW. Upon spawning, the sperm solution was diluted with 500 µL of AFSW and kept on ice. Each female was left to spawn naturally in a Petri dish with 5 mL of AFSW. Approximately 30-50 eggs were transferred to 500 µL of AFSW and fertilized with 10 µL of the sperm solution. Five minutes post-fertilization (pf), excess sperm was removed by moving the eggs to a new dish with 500 µL of AFSW. Eggs were monitored under the stereomicroscope (Leica M165 FC) and photographed at 15 min, 30 min, 1 h, and 3 h pf. The number of fertilized eggs was manually counted from the photographs taken at 30 min pf mark.

## Results

### Overall morphology

We collected specimens along the Ryukyu Archipelago using a hand-held plankton net. Specimens collected from Kin Bay in Okinawa Island (Okinawa lab strain) were cultured in the laboratory alongside the Osaka lab strain (Fig. 1; Material and Methods).

The overall morphologies of the Okinawa lab strain are congruent with previously described *O. dioica* characteristics (Fol 1872; Gorsky and Castellani 2017). It is the only known dioecious appendicularian species; the sexes of Okinawan individuals become visibly distinguishable towards the end of the lifecycle as the gonads at the rear of the body fill with sperm or eggs (Fig. 2A). Viewed dorsally, there are two spindle-shaped subchordal cells to the right of the notochord, generally located in the distal half of the tail (Fig. 2B), with the spacing of the two subchordal cells varying between individuals. The mouth is almost circular with upper and lower lips and there are two spherical buccal glands nearby. The esophagus connects the mouth to the left lobe of the stomach, which then leads to the right lobe. There are round, ciliated spiracles on either side of the rectum that are connected to the pharynx. The U-shaped endostyle is situated ventrally to the pharynx. The brain is present in the anterodorsal region near the pharynx (Fig. 2C, D). According to these morphological criteria, we concluded that the specimens collected in Okinawa were *O. dioica*.

### 18S phylogeny

To compare the Ryukyu specimens to other appendicularian species, we estimated phylogenetic trees of 18S ribosomal rDNA sequences using maximum likelihood (ML) and Bayesian methods. We combined publicly available data from GenBank with 18S rDNA sequences retrieved from the genome sequences of our own and collaborators’ specimens. The resulting trees show that all *O. dioica* specimens form a distinct cluster from other appendicularian species (Fig. 3; ultrabootstrap support,100%; posterior probability, 1.00).

**Fig. 3.**
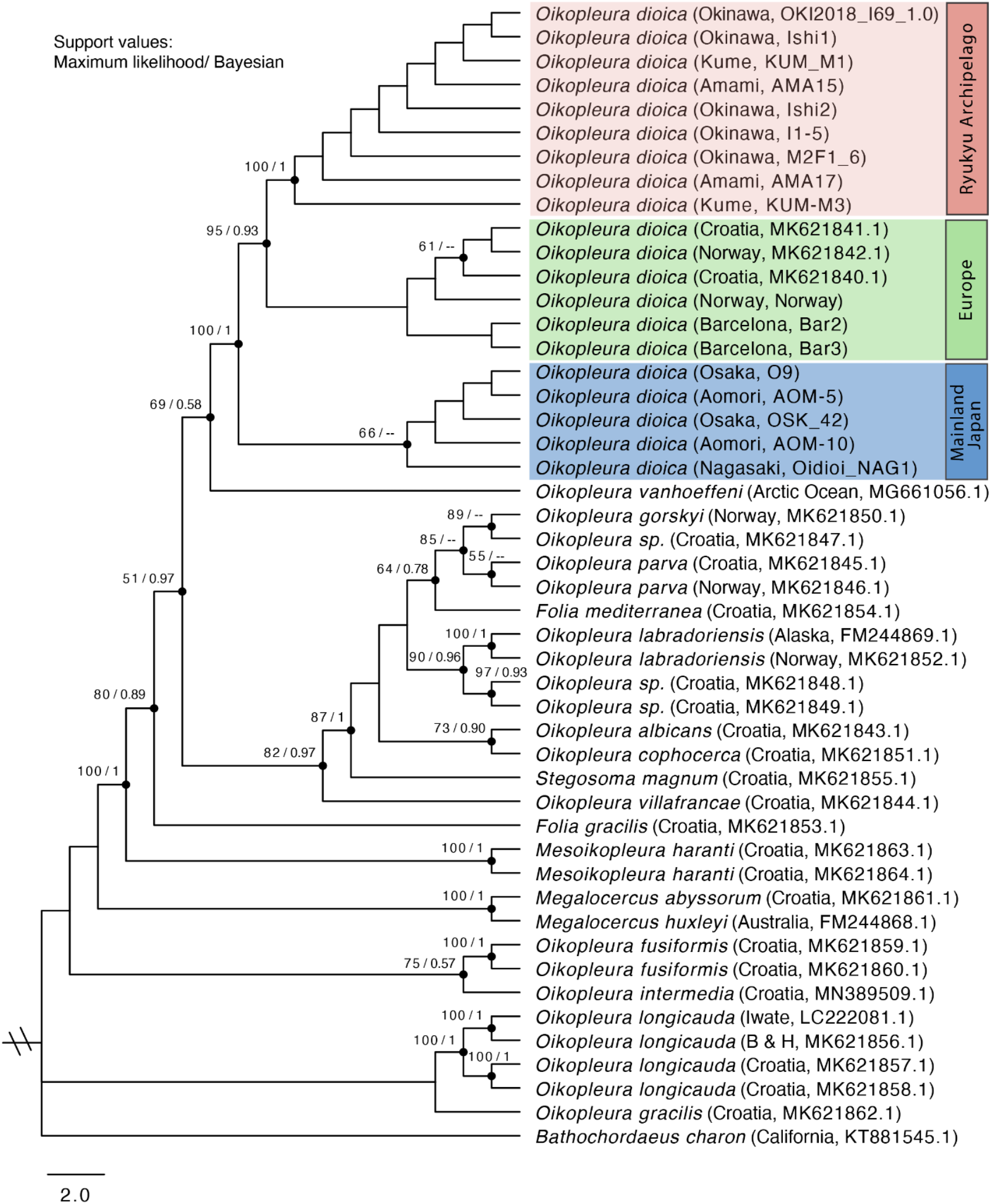
Maximum likelihood phylogenetic tree estimated using ribosomal 18S DNA sequences. Each entry shows the species name, geographical origin, and sample ID. The support values of nodes with >50% bootstrap support and >0.5 posterior probability are shown. Branch support represents maximum likelihood ultrafast bootstrap value (1,000 replicates)/posterior probability (Bayesian inference). Branch length is proportional to the size of the clade. Polytomies in the Bayesian tree are treated as supporting a clade

However, within the *O. dioica* cluster, it appears that the Ryukyu, mainland Japanese and European specimens each form distinct clades. Although the monophyly of the Ryukyu clade is supported by both ML and Bayesian trees, they present conflicting topologies with respect to its placement within *O. dioica* (Fig. 3; Supplementary Figure S1). We reasoned that since *O. dioica* 18S rDNA sequences are highly conserved, with pairwise sequence identities ranging from 99.8 to 100% – one to three nucleotide changes – it is difficult to resolve the close evolutionary relationships within the *O. dioica* monophyletic group (Table 1).

**Table 1.** Pairwise percent identities for the18S, internal transcribed spacer (ITS), and cytochrome oxidase I (COI) nucleotide and protein sequences. The numbers in parentheses indicate the number of mismatched positions for each pairwise alignment

| Aligned sequences | Localities | Species | 18S | ITS | COI (nucleotide) | COI (protein) |
| --- | --- | --- | --- | --- | --- | --- |
| **OKI2018_I69_1.0** | **Okinawa (Ryukyu Archipelago)** | *Oikopleura dioica* |  |  |  |  |
| KUME-M3 | Kume (Ryukyu Archipelago) | *Oikopleura dioica* | 100 (0) | 99.2 (0) |  |  |
| AMA-15 | Amami (Ryukyu Archipelago) | *Oikopleura dioica* | 100 (0) | 99.4 (0) |  |  |
| O9 | Osaka (Mainland Japan) | *Oikopleura dioica* | 99.8 (3) | 87.0 (41) | 99.2 (4) | 85.5 (214) |
| Oidioi_NAG1 | Nagasaki (Mainland Japan) | *Oikopleura dioica* | 99.8 (3) | 87.2 (40) |  |  |
| AOM-10 | Aomori (Mainland Japan) | *Oikopleura dioica* | 99.8 (3) | 87.2 (40) |  |  |
| Norway | Norway (Europe) | *Oikopleura dioica* | 99.8 (3) | 84.2 (44) | 99 (5) | 83.4 (246) |
| Bar2 | Barcelona (Europe) | *Oikopleura dioica* | 99.9 (2) | 87.0 (45) |  |  |
| **O9** | **Osaka (Mainland Japan)** | *Oikopleura dioica* |  |  |  |  |
| KUME-M3 | Kume (Ryukyu Archipelago) | *Oikopleura dioica* | 99.8 (3) | 87.0 (41) |  |  |
| AMA-15 | Amami (Ryukyu Archipelago) | *Oikopleura dioica* | 99.8 (3) | 87.0 (41) |  |  |
| Oidioi_NAG1 | Nagasaki<br>(Mainland Japan) | *Oikopleura dioica* | 100 (0) | 99.7 (0) |  |  |
| AOM-10 | Aomori<br>(Mainland Japan) | *Oikopleura dioica* | 100 (0) | 99.7 (0) |  |  |
| Norway | Norway (Europe) | *Oikopleura dioica* | 99.9 (2) | 93.7 (11) | 99.2 (4) | 85.4 (214) |
| Bar2 | Barcelona (Europe) | *Oikopleura dioica* | 99.9 (1) | 97.0 (10) |  |  |
| **OdB3** | **Norway (Europe)** | *Oikopleura dioica* |  |  |  |  |
| KUME-M3 | Kume (Ryukyu<br>Archipelago) | *Oikopleura dioica* | 99.8 (3) | 84.1 (39) |  |  |
| AMA-15 | Amami (Ryukyu<br>Archipelago) | *Oikopleura dioica* | 99.8 (3) | 84.0 (40) |  |  |
| Oidioi_NAG1 | Nagasaki<br>(Mainland Japan) | *Oikopleura dioica* | 99.9 (2) | 93.7 (11) |  |  |
| AOM-10 | Aomori<br>(Mainland Japan) | *Oikopleura dioica* | 99.9 (2) | 93.7 (11) |  |  |
| Bar2 | Barcelona (Europe) | *Oikopleura dioica* | 99.9 (1) | 98.5 (1) |  |  |
| ***C. intestinalis***<br>**(Roscoff)** | **Roscoff, France** | *Ciona intestinalis* |  |  |  |  |
| *C. robusta*<br>(HT) | Miyagi, Japan | *Ciona robusta*<br>(*Ciona intestinalis*<br>Type A) | 99.7 (0) | 93.1 (27) |  |  |

### ITS phylogeny

To avoid relying on a single molecular marker and improve phylogenetic resolution, we constructed trees using the less conserved, non-coding, internal transcribed spacer (ITS) region of ribosomal loci (Fig. 4). This analysis also generated three highly supported clades within the *O. dioica* monophyletic group. ITS trees indicate that the mainland Japanese and European clades are most closely related, whereas the Ryukyu clade forms a more distant, highly supported lineage. The higher sequence divergence of ITS regions compared with the 18S rDNA genes enables phylogenetic reconstruction at a higher resolution (pairwise sequence identities vary from 84.0% for Amami (AMA-15) vs. Norway (Norway) to 99.7% for Osaka (O9) vs. Nagasaki (Oidioi_NAG1), Osaka (O9) vs. Aomori (AOM-10). In comparison, the sequence identity between *Ciona intestinalis* and the congeneric *Ciona robusta* is 93.1% (Table 1). Although the Ryukyu clade is highly supported in the 18S and ITS trees, the identity of the basal group is ambiguous (Fig. 4, Supplementary Figure S1).

**Fig. 4.**
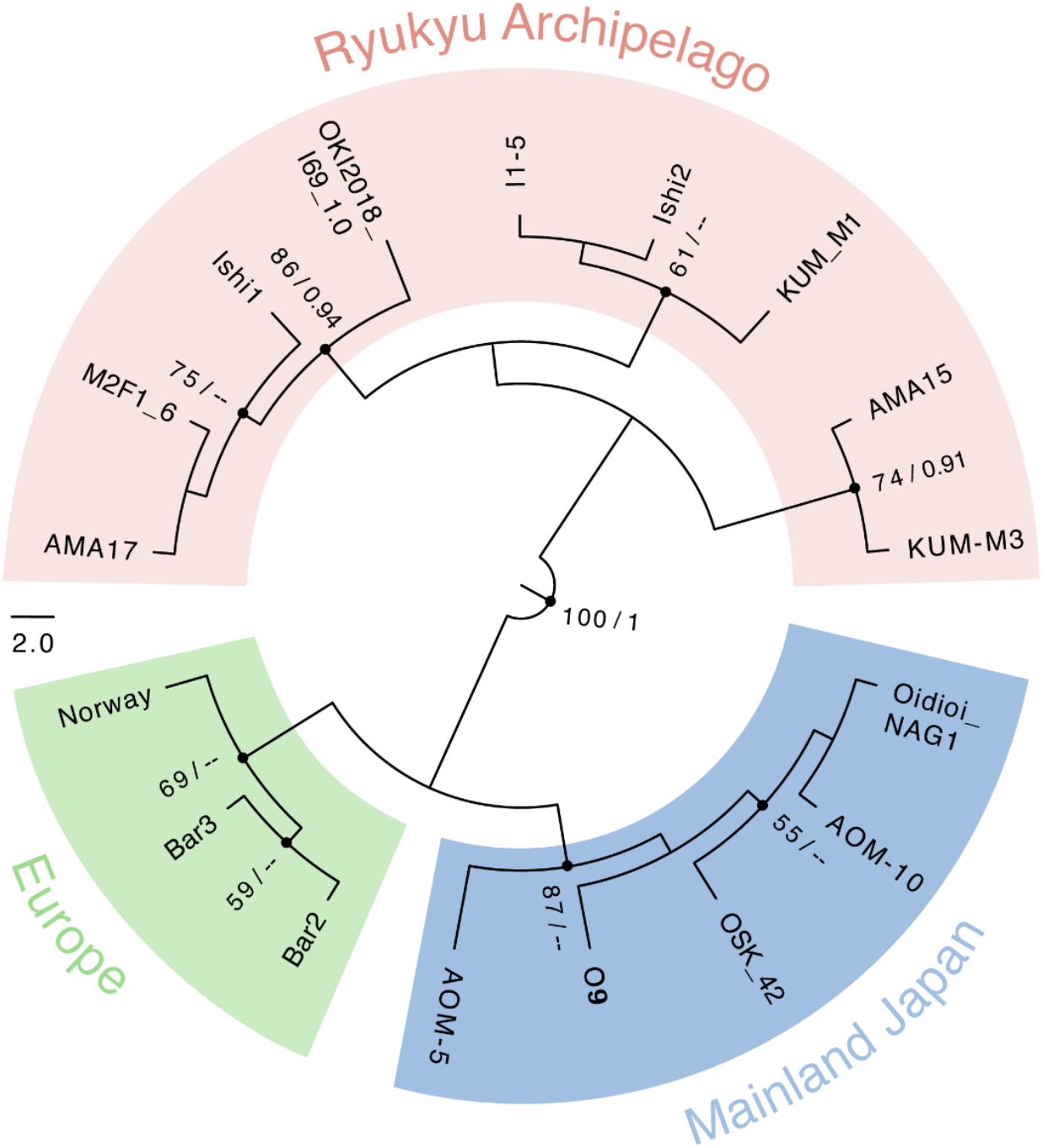
Maximum likelihood phylogenetic tree estimated for *O. dioica* internal transcribed spacer sequences. Support values for nodes with bootstrap support >50% and posterior probability >0.5 are shown. Branch support indicates maximum likelihood ultrafast bootstrap value (1,000 replicates) / posterior probability (Bayesian inference). Branch length is proportional to the size of the clade. Polytomies in the Bayesian tree are treated as supporting a clade

### COI phylogeny

To investigate further genetic differences between the three lineages, and to provide reference data for future studies, we searched the available transcriptomic data for mitochondrial cytochrome oxidase I (COI) sequences in the laboratory strain of each lineage. There are too few samples to construct phylogenetic trees, but alignments show pairwise nucleotide sequence identities between 83.4% and 85.5% (Table 1). Okinawa (OKI2018_I69_1.0) and Norway (Norway) display the greatest pairwise divergence, which is consistent with the ITS comparisons. The translated protein sequences differ by only four or five amino acids in the C-terminal region and most changes occur among the hydrophobic amino acids Val-Leu-Met-Ile and Ala-Thr-Ser.

### Morphological comparisons of the oikoplastic epithelium (OE)

Since our phylogenetic analyses revealed three distinct clades within the *O. dioica* monophyletic group, we decided to examine the morphologies of the oikoplastic epithelium (OE) in greater detail. The OE is a monolayer trunk epithelium responsible for synthesizing the cellulose house that surrounds the animals. The position, number, and shape of OE are considered to be specific to different appendicularian species (Spada et al. 2001; Flood 2005). Cells are arranged in a bilaterally symmetric fashion (Ganot and Thompson 2002); therefore, we examined one side of the OE nuclei patterning in each specimen. We compared the OE nuclear pattern of Okinawa, Osaka, and Barcelona laboratory strains.

We found remarkable conservation in the number of nuclei and the spatial arrangement of the two most prominent domains of the OE (Fol and Eisen) between Okinawa, Osaka, and Barcelona lab strains (Fig. 5, Table 2). For all three lab strains, there are small specimen-to-specimen variations in the numbers of nuclei in the Chain of Pearls (CP) and Nasse (N) regions; these variabilities were also observed in *O. dioica* specimens from Osaka and Norway (Flood 2005; Kishi et al. 2017). Although the first row of the Posterior Fol (PF1) in Osaka was reported to have 10 cells (Kishi et al. 2017), our counts revealed 11 cells for Okinawa (n = 17), Osaka (n = 15), and Barcelona (n = 5) laboratory strains. Nuclear count details are shown in Supplementary Table S2.

**Table 2.** Summary of average counts of the oikoplastic epithelium nuclei in the Fol and the Eisen domains from the Okinawa, Osaka, and Barcelona laboratory strains. The number of nuclei were manually counted using the nomenclature described by Kishi et al. 2017. n = the number of specimens examined

| Domain | Field | Okinawa | Osaka | Barcelona | Kishi et al. 2017 (Osaka) |
| --- | --- | --- | --- | --- | --- |
| Fol | Anterior Crescent (AC) row 1 | 4 | 4 | 4 | 4 |
| Fol | Anterior Crescent (AC) row 2 | 5 | 5 | 5 | 5 |
| Fol | Anterior Crescent (AC) row 3 | 6 | 6 | 6 | 6 |
| Fol | Anterior Crescent (AC) row 4 | 3 | 3 | 3 | 3 |
| Fol | Anterior Elongate (AE) row 1 | 8 | 8 | 8 | 8 |
| Fol | Anterior Elongate (AE) row 2 | 9 | 9 | 9 | 9 |
| Fol | Anterior Elongate (AE) row 3 | 8 | 8 | 8 | 8 |
| Fol | Anterior Elongate (AE) row 4 | 7 | 7 | 7 | 7 |
| Fol | Anterior Elongate (AE) row 5 | 7 | 7 | 7 | 7 |
| Fol | Giant Fol (GF) | 7 | 7 | 7 | 7 |
| Fol | Nasse (N) row 1 | 25 | 26 | 26 | 27 |
| Fol | Nasse (N) row 2 | 25 | 26 | 26 | 27 |
| Fol | Nasse (N) row 3 | 25 | 26 | 26 | 27 |
| Fol | Posterior Fol (PF) row 1 | 11 | 11 | 11 | 10 |
| Fol | Posterior Fol (PF) row 2 | 6 | 6 | 6 | 6 |
| Fol | Posterior Fol (PF) row 3 | 6 | 6 | 6 | 6 |
| Fol | Posterior Fol (PF) row 4 | 5 | 5 | 5 | 5 |
| Fol | Posterior Fol (PF) row 5 | 7 | 7 | 7 | 7 |
| Fol | Posterior Fol (PF) row 6 | 7 | 7 | 7 | 7 |
| Fol | Posterior Fol (PF) row 7 | 8 | 8 | 8 | 8 |
| Fol | Posterior Fol (PF) row 8 | 7 | 7 | 7 | 7 |
| Fol | Posterior Fol (PF) row 9 | 11 | 11 | 11 | 11 |
|  | **Sample size (n)** | 17 | 15 | 5 | 3 |
| Eisen | Eisen Giant cells (E) | 7 | 7 | 7 | 7 |
| Eisen | Eisen's yard (EY) | 20 | 20 | 20 | 20 |
| Eisen | Chain of Pearl (CP) | 13 | 12 | 9 | 11 |
|  | **Sample size (n)** | 16 | 15 | 3 | 3 |

**Fig. 5.**
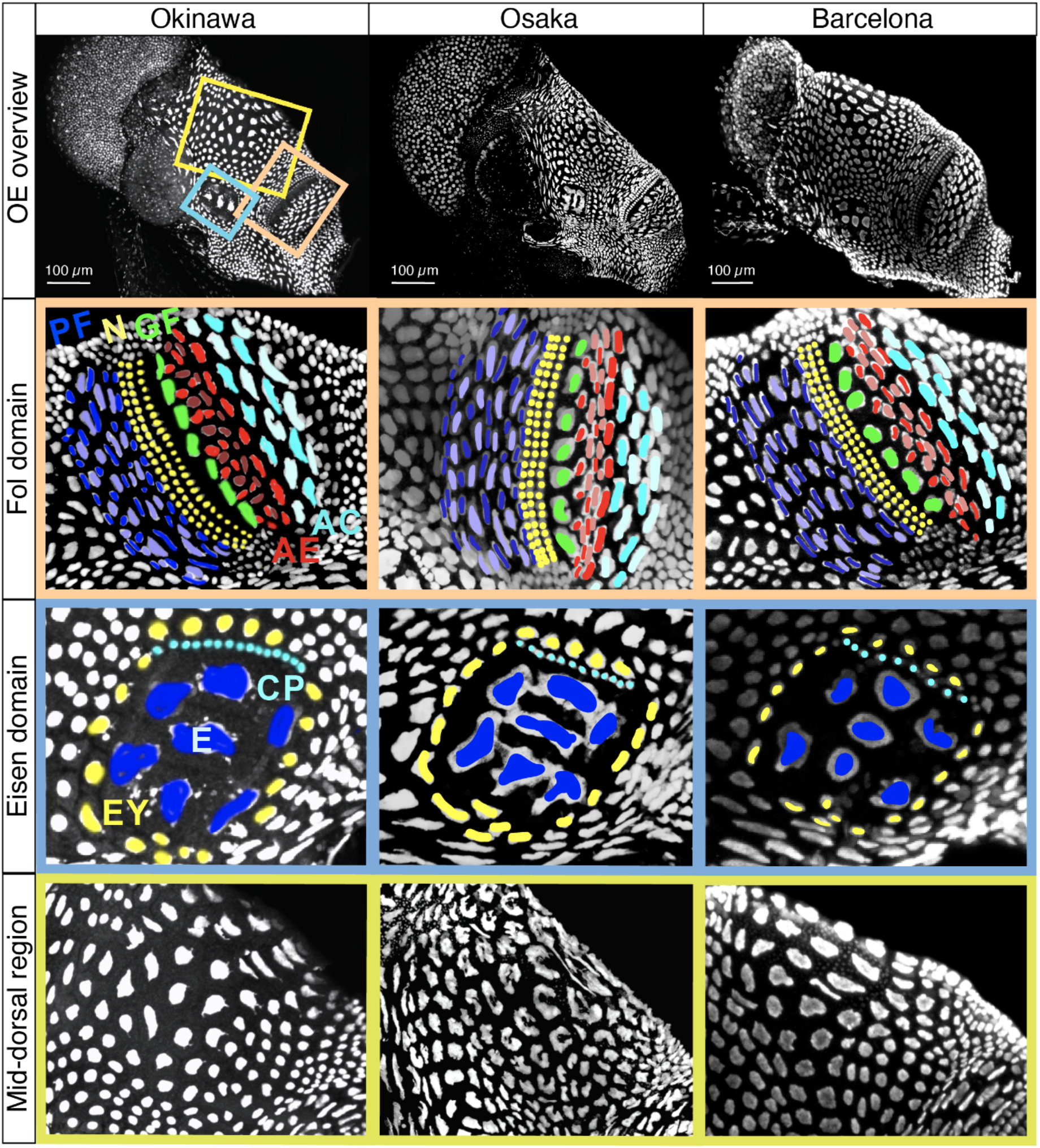
Comparison of the nuclear pattern of the oikoplastic epithelium of Okinawa (left), Osaka (middle), and Barcelona (right) *O. dioica* lab strains. From the top row; the overview of the oikoplastic epithelium, close-up views of the Fol domain (orange), the Eisen domain (blue), and the mid-dorsal regions (yellow). The Fol domain is organized by: Anterior Crescent (AC), Anterior Elongate (AE), Giant Fol (GF), Nasse (N), and Posterior Fol (PF). The Eisen domain is organized by Eisen Giant cells (E), Eisen’s yard (EY), and Chain of Pearl (CP)

The number and the arrangement of the OE nuclei are highly conserved among the Okinawa, Osaka, and Barcelona individuals; however, closer examination revealed noticeable differences in the shape of nuclei in the mid-dorsal regions (Fig. 5). For Osaka animals, the nuclei are crescent-shaped compared with the roughly round shapes of the Okinawa and Barcelona strains. This observation was consistent throughout all the specimens examined during this study (Supplementary Figure S2).

### Trunk/tail ratios and egg diameters

As DNA sequence analysis and visualization of the OE nuclei require instruments that are not yet readily available outside the laboratory, we sought additional morphological markers that are readily accessible in the field. We compared the trunk-to-tail ratios and the average egg diameters for the Okinawa, Osaka, and Barcelona laboratory strains (Fig. 6). There is a statistically significant difference in the trunk-to-tail ratios for Okinawa (mean ± 95% CI of 0.27 ± 0.007, n = 53), Barcelona (0.26 ± 0.006, n = 50), and Osaka strains (0.31 ± 0.004, n = 52; one-way ANOVA, F(2,152) = 68.01, p = 2.2 e^−16^) with overlapping values (Fig. 6A). There is a statistically significant difference in the average egg diameter of Okinawa (mean ± 95% CI of 80 ± 0.0012 µm, n = 20), Osaka (96 ± 0.0015 µm, n = 21) and Barcelona females (96 ± 0.0039 µm, n = 21; one-way ANOVA, F(2,59) = 63.91; p = 1.7e^−15^). Measurements on Okinawa and Osaka egg diameters show no overlapping values (Fig. 6B). Details on the trunk-to-tail ratios and the egg diameter measurements are shown in Supplementary Tables S3 and S4.

**Fig. 6.**
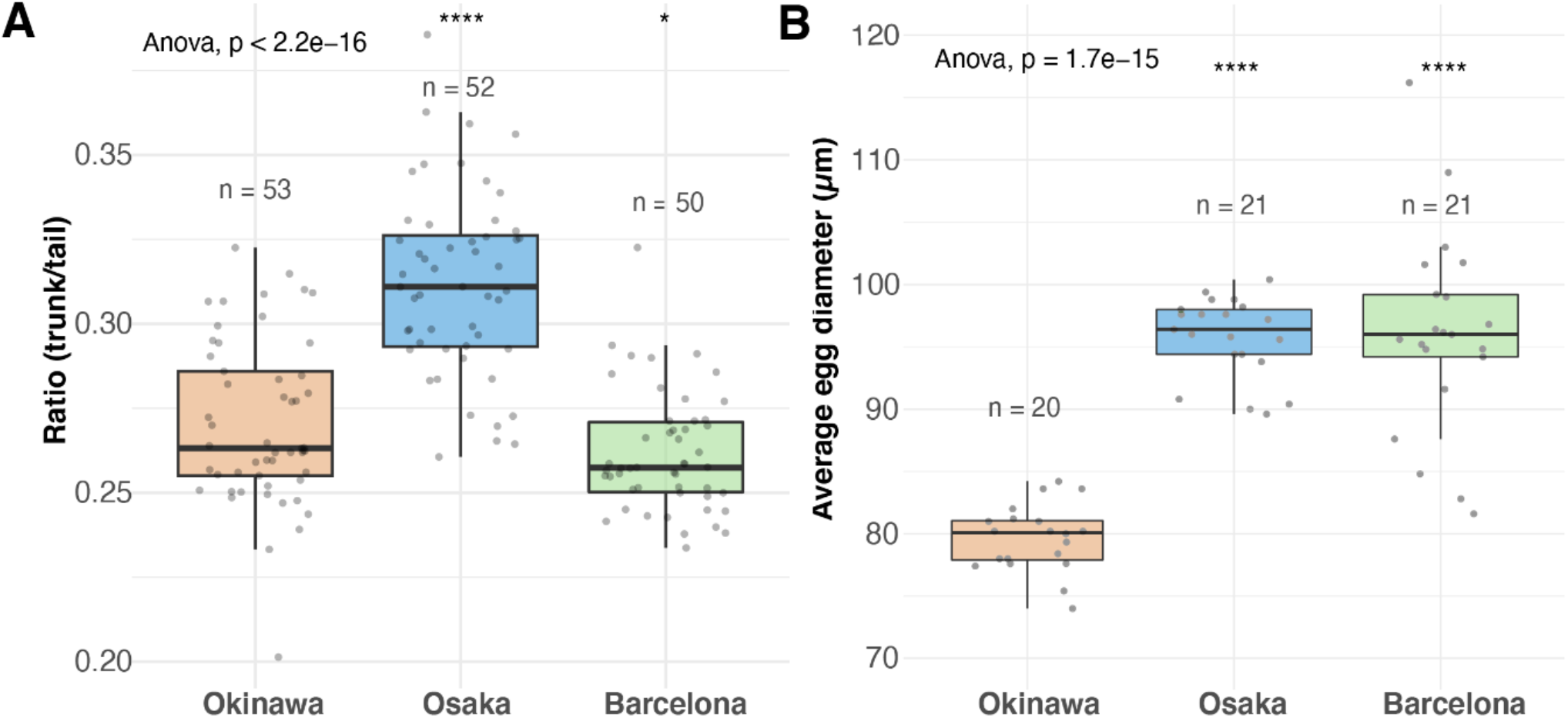
Comparisons of the (A) trunk to tail ratio and the (B) average egg diameter per female individual for the Okinawa, Osaka, and Barcelona laboratory strains. p <= 0.0001 (****), p <= 0.05 (*)

### Gamete compatibility

Sixty Okinawa and Osaka *O. dioica* individuals were selected for *in vitro* fertilization (Fig. 7). 158 out of 179 (88.2%) eggs from Okinawa females and 194 out of 197 (98.5%) eggs from Osaka females were successfully fertilized in the positive control pairs. In contrast, none of the Okinawa-Osaka pairings led to fertilization: eggs inseminated with sperm from non-native males showed no signs of development 3 hours post-fertilization, which is the approximate time to hatching under normal conditions at 20 °C. Individual measurements of the crossing experiment are shown in Supplementary Table S5. Therefore, the Okinawa and Osaka strains are reproductively incompatible.

**Fig. 7.**
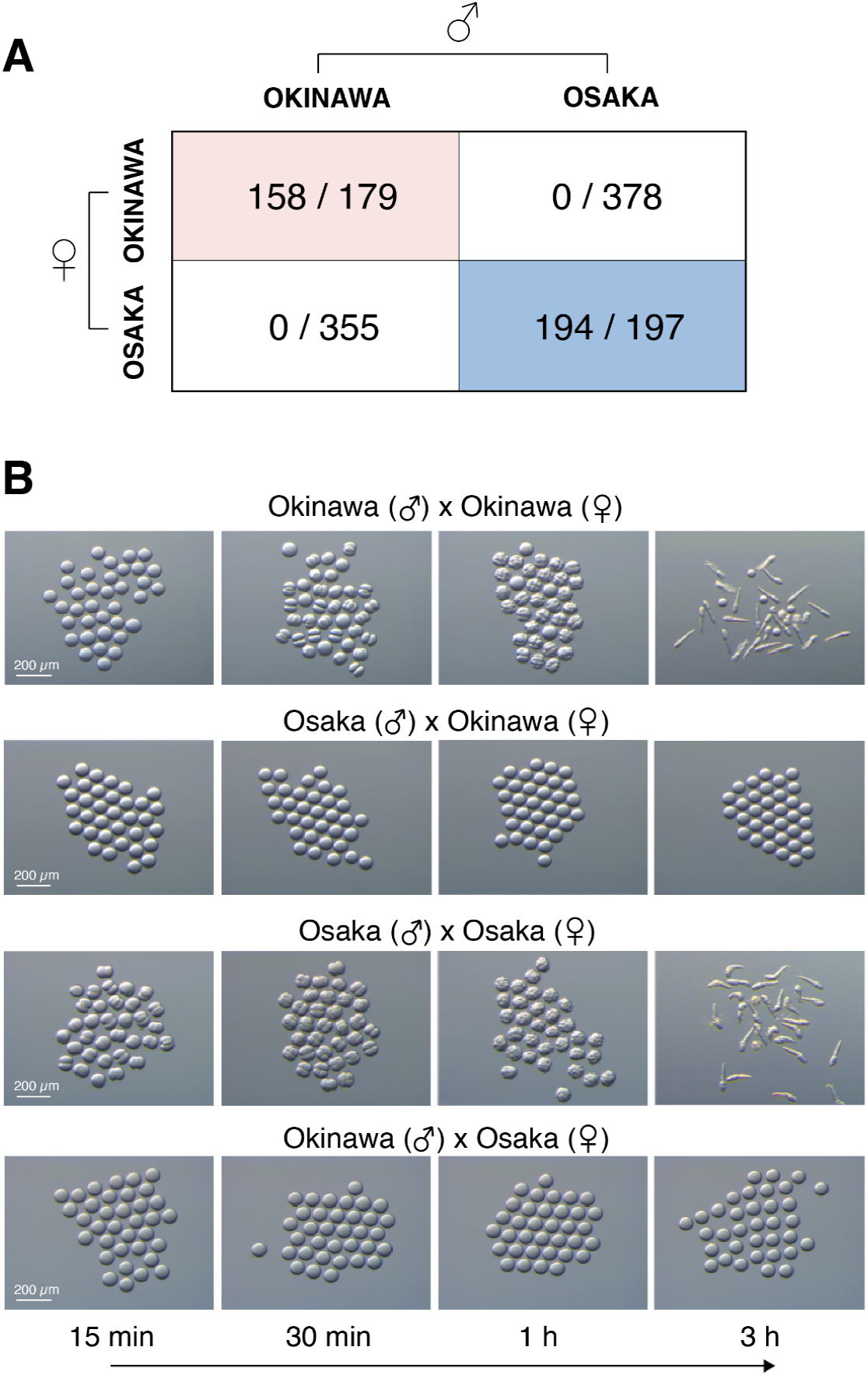
Crossing experiments for Okinawa and Osaka laboratory strains. (A) Total number of eggs successfully fertilized for the Okinawa and Osaka mating pairs. (B) Photographs documenting development after crossing Okinawa and Osaka laboratory strains. Photographs were taken after 15 min, 30 min, 1h, and 3h post-fertilization attempt

## Discussion

Here we presented evidence for cryptic speciation of *O. dioica*. We assessed three orthogonal pieces of evidence: phylogeny (Cracraft 1983), morphology (Cronquist 1978), and reproductive compatibility (Mayr 1942). Although the Ryukyu, mainland Japanese, and European specimens exhibit remarkable conservation in morphology, our phylogenetic analyses suggest they comprise three genetically distinct lineages that are currently considered conspecific. It is possible, indeed likely, that the number of lineages will increase as samples from additional geographical locations are analysed in future studies.

### Phylogeny

18S rDNA gene is a useful marker for distinguishing appendicularian species; however, it is too conserved to discriminate closely related *O. dioica* lineages (Fig 3). The non-coding ITS region is useful for distinguishing recently diverged taxa within the *O. dioica* monophyletic group (Fig 4). We therefore suggest ITS region as a candidate marker for *O. dioica* lineage identification and delimitation. The ITS region has been widely used in DNA barcoding analyses in many taxa including algae, protists, and animals and has been proposed as the standard DNA barcode for fungi and seed plants (Schlötterer et al. 1994; Yao et al. 2010; Wang et al. 2015b).

In addition to the three global lineages of *O. dioica*, there is local diversity within the Ryukyu clade (Fig. 4): there are three groups comprising (i) two individuals from Amami (AMA 15) and Kume islands (KUM-M3), (ii) another Amami individual (AMA 17) that clusters with specimens from Okinawa island (OKI2018_I69_1.0, Ishi1, and M2F1_6), and (iii) a Kume individual (KUM_M1) that clusters with two other Okinawa individuals (I1-5 and Ishi2). Despite this diversity, the placements suggest that specimens from Okinawa, Kume, and Amami islands form a single, continuous Ryukyu *O. dioica* population. Further examination of local samples will provide insights into the population structure and the distribution of the Ryukyu *O. dioica* clade and its boundaries with neighboring populations.

### Morphology

The overall morphologies are hardly distinguishable between the Okinawan, Osaka, and Barcelona *O. dioica* specimens. Closer examination in the trunk-to-tail ratios and egg diameters reveal subtle quantitative differences. However, it is important to note that morphological measurements were taken from the specimens grown in laboratories. The size of animals including gonad length and egg diameters may vary depending on culture conditions, such as water quality and food availability.

The oikoplastic epithelium (OE), which is responsible for creating the house structure in appendicularians species, is highly conserved between the three genetic *O. dioica* lineages. Only in the mid-dorsal region is it possible to observe distinct crescent-shaped nuclei in the Osaka specimens (Fig. 5). A similar observation was reported among *Oikopleura longicauda*, in which prominent differences in size and shape of the OE nuclei were found between specimens collected from different geographical regions (Flood 2005).

Many larger appendicularian species (trunk length > 1.4 mm), such as *Oikopleura labradoriensis, O. vanhoeffeni*, and *O. villafrancae*, exhibit more complex and arborized patterns of OE nuclei compared with smaller appendicularians such as *O. dioica, O. gorskyi, O. parva*, and *O. longicauda* (Flood 2005). These differences may reflect the changes in size and ploidy of OE cells. In *O. dioica*, OE cells stop mitotic division prior to metamorphosis and continue their growth by increasing the nuclear DNA content and size of nuclei. The nuclear morphologies change with the ploidy of OE cells as animals develop (Ganot and Thompson 2002). Therefore, despite our efforts to keep consistent staging throughout sample collection, it is possible that the observed differences in OE nuclei shape could arise from differences in the developmental stages of individual specimens. The primary observation is that the number and arrangement of OE cells are highly conserved among genetically distinct lineages of *O. dioica*. Overall, the morphologies of the three *O. dioica* lineages are indistinguishable at a superficial level. Further studies are needed to evaluate the possible existence of morphological markers that can reliably distinguish *O. dioica* lineages.

### Reproductive isolation

Finally, we investigated the fertilization success of Okinawa and Osaka lab strains. Genetically distinct lineages of *O. dioica* from the Okinawa and Osaka lab strains do not interbreed under experimental conditions (Fig. 7). Reciprocal fertilization trials using individuals from these two strains always resulted in failure, suggesting strong prezygotic mechanisms against fertilization.

The intensity of this reproductive barrier between two relatively close geographical locations initially surprised us. It could be explained by the Kuroshio current - the subtropical western boundary current in the North Pacific that flows northwestwards along Okinawa Island and northeastwards along mainland Japan. The warm Kuroshio current transports many tropical and subtropical species to the north (Saito 2019). However, it can also act as a dispersal barrier, promoting lineage diversification in marine organisms (Kojima et al. 2000). Although appendicularians are present in the Kuroshio ecosystem (Saito 2019), *O. dioica* in particular is an inlet species and is most commonly found in bays and harbors (Tomita et al. 2003; Hidaka 2008). Therefore, even if a small population of *O. dioica* exists in the Kuroshio, the fast and strong meandering current at the Tokara gap (Fig. 1) may act as a potential dispersal barrier, limiting gene flow (Ogoh and Ohmiya 2005; Liu et al. 2008; Yamazaki et al. 2017) between Ryukyu and mainland Japanese *O. dioica*.

### Implications and next steps

*O. dioica* is an important model organism for appendicularian and tunicate biology, partly owing to its global distribution, relative ease of sample collection, and ability to culture long-term in the laboratory. The original molecular comparisons between Norwegian and North Pacific specimens suggested a high level of DNA sequence conservation; however, more recent genomic and transcriptomic comparisons revealed more extensive divergence at the single nucleotide level. This study now confirms the impact of this divergence on the molecular phylogeny of different *O. dioica* specimens.

Studies have reported key developmental features that are not only common across European, Japanese and North American *O. dioica*, but also among chordates. In light of the high level of morphological conservation and similarities in developmental trajectories and pathways, many molecular, cellular and developmental biology discoveries in the different lineages will continue to apply across all *O. dioica*. Indeed, of great interest will be to understand the similarities and differences of distinct lineages, particularly in the context of their native marine environments.

Our work provides the first evidence that *O. dioica* might be hiding multiple cryptic species and establishes the basis for further investigations into the extent of *O. dioica* genetic diversity around the globe. The evidence suggests that at least the Ryukyu specimens should be considered a separate species to European and mainland Japanese *O. dioica*, which might require revision of current *O. dioica* taxonomy. Our findings also raise interesting new questions to be tackled in future studies. What is the extent of geographical distributions of different lineages? Are different lineages sympatric or not? What is the level of genetic diversity within each lineage? In what way does the loss of repair mechanisms impact the evolution of these organisms? Approaches targeting environmental DNA (eDNA) using ITS primers may provide molecular data that help us detect and distinguish *O. dioica* lineages. It is necessary to examine whether practical morphological characteristics could be discovered to aid field identification. We anticipate that the combined approach incorporating morphological and molecular markers will facilitate greater exploration into the diversity and biogeography of these ecologically important animals for ocean’s health.

## Data availability

All sequence data have been deposited in the GenBank database (accession numbers provided in Supplementary Table S1). Raw data of our experiments are provided in the electronic supplementary files.

## Acknowledgment

We are grateful to Hiroki Nishida, Hitoshi Akiyama, Astuo Nishino, Kouhei Hirose, Shouchi Nishino, for sending *Oikopleura* samples to us. We are thankful to Rade Garić for the advice and suggestion on taxonomy and morphological measurements. We thank Shinnosuke Teruya for allowing us to use the Deep Sea water research center on Kume Island. We are also thankful to the Kujukushima Aquarium Umi Kirara staff, especially Hisashi Akiyama, for their effort to identify and collect *O. dioica* for us. We thank Jai Denton for his support and encouragement, especially during the early phase of the project. We thank Shinya Komoto from the OIST Imaging Section for assisting with Oikopleura epithelium imaging. Finally, we thank the Okinawa Institute of Science and Technology Graduate University (OIST) for funding, the DNA Sequencing Section and the Scientific Computing and Data Analysis Section of the Research Support Division at OIST, and the OIST Fieldwork Safety Committee for advice on safe sampling procedures. M.J.M. gratefully acknowledges funding from the Japan Society for the Promotion of Science as a JSPS International Research Fellow (Luscombe Unit, Okinawa Institute of Science and Technology Graduate University)

## Contributions

*Aki Masunaga, Charles Plessy, Michael Mansfield, and Nicholas* Luscombe contributed to the study conception and design. Material preparation was performed by Aki Masunaga, Yongkai Tan, Andrew Liu, Aleksandra Bliznina, and Takeshi Onuma. Data collection was performed by Aki Masunaga, Yongkai Tan, Paolo Barzaghi, Cristian Cañestro and Alfonso Ferrández-Roldán. Data analysis was performed by Aki Masunaga, Tamara Hodgetts, Michael Mansfield, and Charles Plessy. The first draft of the manuscript was written by Aki Masunaga and all authors read and approved the final manuscript.

## Ethics declarations

### Ethics approval

All applicable national and institutional guidelines for the care and use of animals were followed. This article does not contain any studies with human participants. All appendicularian specimen collections were approved by the OIST Fieldwork Safety Committee.

## Supplementary Information

Below is the link to electronic supplementary materials. https://docs.google.com/document/d/1w7n5XWxFGnClaskFR92VMgo8-jExbL3nduMFeL2PY2I/edit

