## Supplementary materials for "The cosmopolitan appendicularian *Oikopleura dioica* reveals hidden genetic diversity around the globe"

**Supplementary Table S1** Sample information and ribosomal DNA locus Genbank Accession numbers.

| Species | 18S GenBank accession number | rDNA locus GenBank accession number | Sample ID | Locality | Reference |
|---|---|---|---|---|---|
| *Oikopleura dioica* | — | OP113812 | OKI2018_I69_1.0 (Okinawa lab strain) | Okinawa, Japan | Masunaga et al. 2022 |
| *Oikopleura dioica* | — | OP113810 | Ishi2 (Okinawa lab strain) | Okinawa, Japan | Masunaga et al. 2022 |
| *Oikopleura dioica* | — | OP113801 | I1-5 (Okinawa lab strain) | Okinawa, Japan | Masunaga et al. 2022 |
| *Oikopleura dioica* | — | OP113811 | M2F1_6 (Okinawa lab strain) | Okinawa, Japan | Masunaga et al. 2022 |
| *Oikopleura dioica* | — | OP113803 | KUM_M1 (wild animal) | Kume, Japan | Masunaga et al. 2022 |
| *Oikopleura dioica* | — | OP113799 | AMA15 (wild animal) | Amami, Japan | Masunaga et al. 2022 |
| *Oikopleura dioica* | — | OP113800 | AMA17 (wild animal) | Amami, Japan | Masunaga et al. 2022 |
| *Oikopleura dioica* | — | OP113802 | KUM-M3 | Kume, Japan | Masunaga |

| | | | | | |
|---|---|---|---|---|---|
| | | | (wild animal) | | et al. 2022 |
| *Oikopleura dioica* | — | OP113813 | Norway (Norway lab strain) | Norway | Masunaga et al. 2022 |
| *Oikopleura dioica* | — | OP113814 | Bar2 (Barcelona lab strain) | Barcelona, Spain | Masunaga et al. 2022 |
| *Oikopleura dioica* | — | OP113815 | Bar3 (Barcelona lab strain) | Barcelona, Spain | Masunaga et al. 2022 |
| *Oikopleura dioica* | — | OP113806 | O9 (Osaka lab strain) | Hyogo, Japan | Masunaga et al. 2022 |
| *Oikopleura dioica* | — | OP113807 | AOM-5 (wild animal) | Aomori, Japan | Masunaga et al. 2022 |
| *Oikopleura dioica* | — | OP113805 | OSK_42 (wild animal) | Hyogo, Japan | Masunaga et al. 2022 |
| *Oikopleura dioica* | — | OP113808 | AOM-10 (wild animal) | Aomori, Japan | Masunaga et al. 2022 |
| *Oikopleura dioica* | — | OP113804 | Oidioi_NAG1 (wild animal) | Nagasaki, Japan | Masunaga et al. 2022 |
| *Oikopleura dioica* | MK621840.1 | — | — | Croatia | Garic et al. Unpublished |
| *Oikopleura dioica* | MK621841.1 | — | — | Croatia | Garic et al. Unpublished |
| *Oikopleura dioica* | MK621842.1 | — | — | Norway | Garic et al. Unpublished |
| *Oikopleura vanhoeffeni* | MG661056.1 | — | — | Arctic ocean, USA | Questel et al. Unpublished |
| *Oikopleura gorskyi* | MK621850.1 | — | — | Norway | Garic et al. Unpublished |
| *Oikopleura sp* | MK621847.1 | — | — | Croatia | Garic et al. Unpublished |
| *Oikopleura parva* | MK621845.1 | — | — | Croatia | Garic et al. Unpublished |

| | | | | | |
|---|---|---|---|---|---|
| *Oikopleura parva* | MK621846.1 | — | — | Norway | Garic et al. Unpublished |
| *Folia mediterranea* | MK621854.1 | — | — | Croatia | Garic et al. Unpublished |
| *Oikopleura albicans* | MK621843.1 | — | — | Croatia | Garic et al. Unpublished |
| *Oikopleura cophocerca* | MK621851.1 | — | — | Croatia | Garic et al. Unpublished |
| *Oikopleura labradoriensis* | FM244869.1 | — | — | Alaska, USA | Tsagkogeorga et al. 2009 |
| *Oikopleura labradoriensis* | MK621852.1 | — | — | Norway | Garic et al. Unpublished |
| *Oikopleura sp.* | MK621848.1 | — | — | Croatia | Garic et al. Unpublished |
| *Oikopleura sp.* | MK621849.1 | — | — | Croatia | Garic et al. Unpublished |
| *Stegosoma magnum* | MK621855.1 | — | — | Croatia | Garic et al. Unpublished |
| *Oikopleura villafrancae* | MK621844.1 | — | — | Croatia | Garic et al. Unpublished |
| *Folia gracilis* | MK621853.1 | — | — | Croatia | Garic et al. Unpublished |
| *Mesoikopleura haranti* | MK621863.1 | — | — | Croatia | Garic et al. Unpublished |
| *Mesoikopleura haranti* | MK621864.1 | — | — | Croatia | Garic et al. Unpublished |
| *Megalocercus abyssorum* | MK621861.1 | — | — | Croatia | Garic et al. Unpublished |
| *Megalocercus huxleyi* | FM244868.1 | — | — | Australia | Tsagkogeorga et al. 2009 |

| | | | | | |
|---|---|---|---|---|---|
| *Oikopleura fusiformis* | MK621859.1 | — | — | Croatia | Garic et al. Unpublished |
| *Oikopleura fusiformis* | MK621860.1 | — | — | Croatia | Garic et al. Unpublished |
| *Oikopleura intermedia* | MN389509.1 | — | — | Croatia | Garic et al. Unpublished |
| *Oikopleura longicauda* | LC222081.1 | — | — | Iwate, Japan | Sakaguchi et al. 2017 |
| *Oikopleura longicauda* | MK621856.1 | — | — | Bosnia and Herzegovina | Garic et al. Unpublished |
| *Oikopleura longicauda* | MK621857.1 | — | — | Croatia | Garic et al. Unpublished |
| *Oikopleura longicauda* | MK621858.1 | — | — | Croatia | Garic et al. Unpublished |
| *Oikopleura gracilis* | MK621862.1 | — | — | Croatia | Garic et al. Unpublished |
| *Bathochordaeus charon* | KT881545.1 | — | — | California, USA | Sherlock et al. 2015 |
| *Ciona intestinalis* | BNKA01000151.1_21137-22948 | — | — | Roscoff, France | Satou et al. 2021 |
| *Ciona robusta* (*Ciona intestinalis Type A*) | BJTB01000029.1_8553-10364 | — | — | Miyagi, Japan | Satou et al. 2019 |

**Supplementary Figure S1** Larvacean 18S phylogenetic tree estimated with maximum likelihood (ML) using (**A**) Treepuzzel and (**B**) PhyML software. *O. dioica* ITS phylogenetic tree estimated with maximum likelihood (ML) using (**C**) Treepuzzel and (**D**) PhyML software. Branch length is proportional to the size of the clade

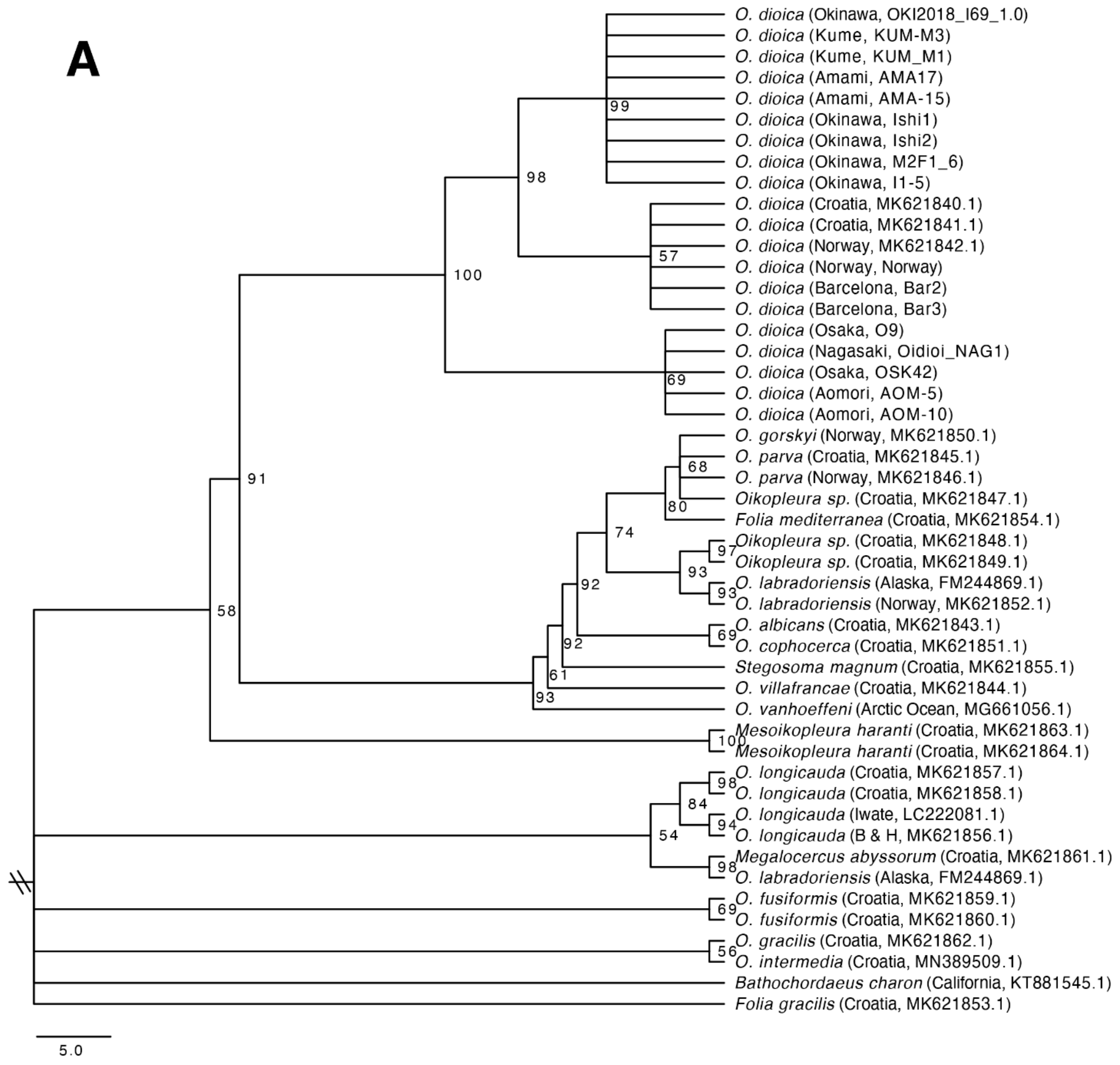

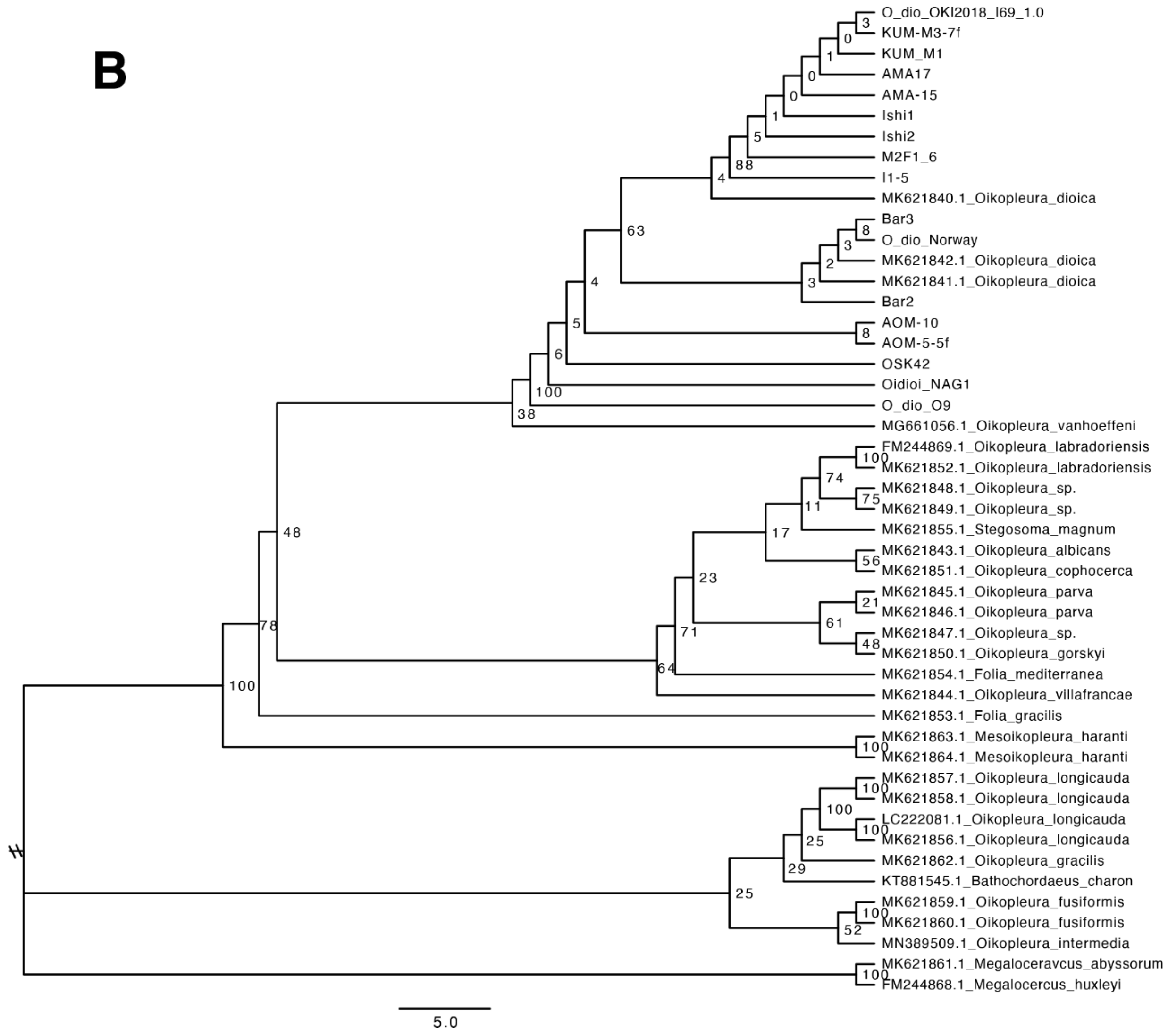
B
O_dio_OKI2018_I69_1.0
KUM-M3-7f
KUM_M1
AMA17
AMA-15
Ishi1
Ishi2
M2F1_6
I1-5
MK621840.1_Oikopleura_dioica
Bar3
O_dio_Norway
MK621842.1_Oikopleura_dioica
MK621841.1_Oikopleura_dioica
Bar2
AOM-10
AOM-5-5f
OSK42
Oidioi_NAG1
O_dio_O9
MG661056.1_Oikopleura_vanhoeffeni
FM244869.1_Oikopleura_labradoriensis
MK621852.1_Oikopleura_labradoriensis
MK621848.1_Oikopleura_sp.
MK621849.1_Oikopleura_sp.
MK621855.1_Stegosoma_magnum
MK621843.1_Oikopleura_albicans
MK621851.1_Oikopleura_cophocerca
MK621845.1_Oikopleura_parva
MK621846.1_Oikopleura_parva
MK621847.1_Oikopleura_sp.
MK621850.1_Oikopleura_gorskyi
MK621854.1_Folia_mediterranea
MK621844.1_Oikopleura_villafrancae
MK621853.1_Folia_gracilis
MK621863.1_Mesoikopleura_haranti
MK621864.1_Mesoikopleura_haranti
MK621857.1_Oikopleura_longicauda
MK621858.1_Oikopleura_longicauda
LC222081.1_Oikopleura_longicauda
MK621856.1_Oikopleura_longicauda
MK621862.1_Oikopleura_gracilis
KT881545.1_Bathochordaeus_charon
MK621859.1_Oikopleura_fusiformis
MK621860.1_Oikopleura_fusiformis
MN389509.1_Oikopleura_intermedia
MK621861.1_Megaloceravcus_abyssorum
FM244868.1_Megalocercus_huxleyi
5.0

**C**

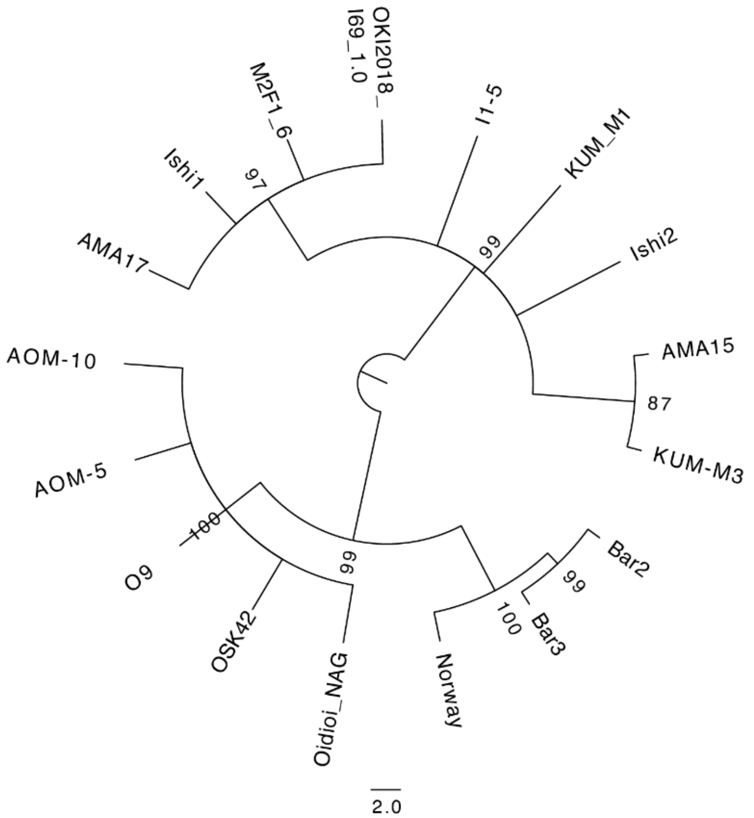

OKI2018_
I69_1.0
M2F1_6
I1-5
KUM_M1
Ishi1
97
Ishi2
99
AMA17
AMA15
87
KUM-M3
AOM-10
AOM-5
100
Bar2
99
O9
99
100
Bar3
OSK42
Oidioi_NAG
Norway
2.0


**D**

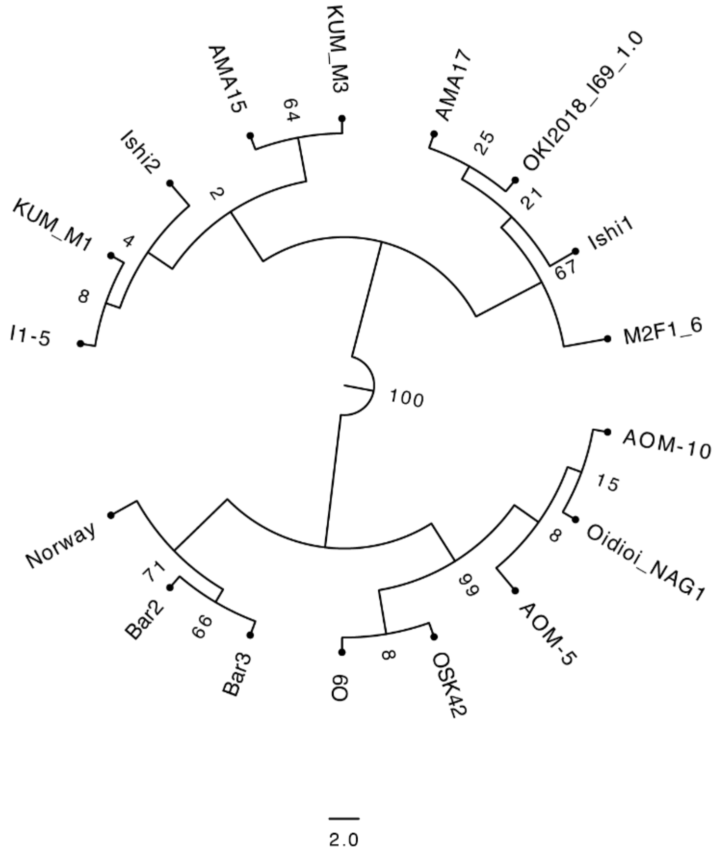

KUM_M3
AMA15
64
AMA17
OKI2018_I69_1.0
25
Ishi2
2
21
KUM_M1
4
Ishi1
67
8
I1-5
M2F1_6
100
AOM-10
15
8
Oidioi_NAG1
Norway
71
99
Bar2
66
AOM-5
Bar3
O9
8
OSK42
2.0

**Supplementary Table S2** Raw counts of the oikoplastic epithelium nuclei in the Fol and the Eisen domains from the Okinawa, Osaka, and Barcelona laboratory strains

| Photo ID | AC1 - AC4 | AE1 - AE5 | GF | N1 - N3 | PF1 - PF10 | EG | EY | CP |
|---|---|---|---|---|---|---|---|---|
| 02092021_ Oki_01 | 4,5,6,3 | 8,9,8,7,7 | 7 | 24,24,24 | 11,6,6,5,7,7,8,-,-,- | 7 | 20 | 13 |
| 02092021_ Oki_02 | 4,5,6,3 | 8,9,8,7,7 | 7 | 25,25,25 | 11,6,6,5,7,7,8,7,11,10 | | | |
| 03182021_ Oki_01 | 4,5,6,3 | 8,9,8,7,7 | 7 | 25,25,25 | 11,6,6,5,7,7,8,7,11,11 | | | 12 |
| 03182021_ Oki_02 | 4,5,6,3 | 8,9,8,7,7 | 7 | 25,25,25 | 11,6,6,5,7,7,9,7,11,10 | | | 12 |
| 03182021_ Oki_03 | 4,5,6,3 | 8,9,8,7,7 | 6 | 25,25,25 | 11,6,6,5,7,7,8,7,11,11 | | | 13 |
| 03182021_ Oki_04 | 4,5,6,3 | 8,9,8,7,7 | | 25,25,25 | 11,6,6,5,7,7,8,7,11,11 | 7 | 20 | 15 |
| 03182021_ Oki_05 | 4,5,6,3 | 8,9,8,7,7 | 7 | 25,25,25 | 11,6,6,5,7,7,8,7,11,10 | 7 | - | 13 |
| 03182021_ Oki_06 | 4,5,6,3 | 8,9,8,7,7 | 7 | 25,25,25 | 11,6,6,5,7,7,8,7,11,12 | 7 | 20 | 12 |
| 04152021_ Oki_01 | 4,5,6,3 | 8,9,8,7,7 | 7 | 23,23,23 | 11,6,6,5,7,7,8,7,11,11 | | | |
| 04152021_ Oki_02 | 3,4,6,3 | 8,9,8,7,7 | 7 | 25,25,25 | 10,6,6,5,7,-,-,-,-,- | | | |
| 04152021_ Oki_03 | 4,5,6,3 | 8,9,8,5,6 | 7 | 25,25,25 | 11,6,6,5,7,7,8,7,11,9 | | | |
| 04152021_ Oki_04 | 4,5,6,3 | 8,9,8,7,7 | 7 | 24,24,24 | 11,6,6,5,7,7,8,7,11,9 | | | |
| 04152021_ Oki_05 | | | 7 | 26,26,26 | 11,6,6,5,7,7,8,7,11,11 | | | |
| 04152021_ Oki_06 | 4,5,6,3 | 7,8,7,5,6 | 6 | 25,25,25 | 11,6,6,5,7,7,8,7,11,7 | | | |
| 04152021_ Oki_07 | 4,5,6,3 | 7,9,8,6,7 | 6 | 25,25,25 | 11,6,6,5,7,7,8,7,11,9 | | | |
| 04152021_ Oki_08 | | | 7 | | 10,6,6,5,7,7,8,7,11,7 | | | |
| 04152021_ Oki_09 | 4,5,6,3 | 8,9,8,7,7 | 7 | 25,25,25 | 11,6,6,5,7,7,8,7,11,8 | | | |
| 04152021_ Oki_10 | | | | | | 7 | 20 | 13 |
| 04152021_ Oki_11 | | | | | | 7 | 20 | 13 |
| 04152021_ Oki_12 | | | | | | 7 | 20 | 13 |
| 04152021_ Oki_13 | | | | | | 7 | 20 | 13 |
| 04152021_ Oki_14 | | | | | | 7 | 20 | 15 |

| | | | | | | | | |
|---|---|---|---|---|---|---|---|---|
| 04152021_ Oki_15 | | | | | | 7 | - | 12 |
| 04152021_ Oki_16 | | | | | | 7 | 20 | 13 |
| 04152021_ Oki_17 | | | | | | 7 | 20 | 12 |
| **Photo ID** | **AC1 - AC4** | **AE1 - AE5** | **GF** | **N1 - N3** | **PF1 - PF10** | **EG** | **EY** | **CP** |
| 09272021_ Osa_01 | 4,5,6,3 | 8,-,-,-,7 | 7 | 25,25,25 | 9,6,6,5,7,7,6,-,-,- | 7 | 20 | 12 |
| 09272021_ Osa_02 | 4,5,6,3 | 7,8,6,7,7 | 7 | 25,25,25 | 11,6,6,5,7,7,6,-,-,- | 7 | 20 | 12 |
| 09272021_ Osa_03 | | | | | | 7 | 20 | 9 |
| 09272021_ Osa_04 | 4,5,6,3 | | | 27,27,27 | | | | |
| 09272021_ Osa_05 | | | | | | 7 | 20 | 12 |
| 09272021_ Osa_06 | 4,5,6,3 | 8,9,7,7,8 | 7 | 27,27,27 | 11,6,6,5,7,78,7,11,- | | | |
| 09272021_ Osa_07 | | | | | | 7 | 20 | 11 |
| 09272021_ Osa_08 | 4,5,6,3 | 8,9,7,6,- | | 27,27,27 | 10,6,6,5,7,78,7,11 | | | |
| 09272021_ Osa_09 | | | | | | 7 | 20 | 11 |
| 09272021_ Osa_10 | 4,5,6,3 | 8,7,-,-,- | | 27,27,27 | 11,6,6,5,7,7,-,-,-,- | | | |
| 09272021_ Osa_11 | | | | | | 7 | 20 | 11 |
| 09272021_ Osa_12 | 4,5,6,3 | 8,9,8,7,7 | | 27,27,27 | 11,6,6,5,7,7,7,7,11 | | | |
| 09272021_ Osa_13 | | | | | | 7 | 21 | 12 |
| 09272021_ Osa_14 | 4,5,6,3 | 8,9,8,7,- | | 27,27,27 | | | | |
| 09272021_ Osa_15 | | | | | | 7 | 21 | 12 |
| 09272021_ Osa_16 | 4,5,6,3 | 8,9,6,-,- | | | 11,6,6,5,7,77,7,-,- | | | |
| 09272021_ Osa_17 | | | | | | 7 | 20 | 12 |
| 09272021_ Osa_18 | 4,5,6,3 | 8,9,-,-,- | | | | | | |
| 09272021_ Osa_19 | 4,5,6,3 | 8,9,8,7,7 | 7 | 27,27,27 | 11,6,6,5,7,7,7,7,11,- | | | |
| 09272021_ Osa_20 | | | | | 11,6,6,5,7,7,7,7,11,- | | | |

| | | | | | | | | |
|---|---|---|---|---|---|---|---|---|
| 09272021_ Osa_21 | 4,5,6,3 | 8,9,-,-,- | | 27,27,27 | 11,6,6,5,7,7,8,7,9,- | | | |
| 09272021_ Osa_22 | 4,5,6,3 | 8,7,8,6,6 | 7 | 27,27,27 | 11,6,6,5,7,7,7,7,11,- | | | |
| 09272021_ Osa_23 | 4,5,6,3 | 8,-,-,-,- | 7 | 26,26,26 | 11,6,6,5,7,7,-,-,-,- | | | |
| 09272021_ Osa_24 | 4,5,6,3 | 8,9,8,7,7 | 7 | 26,26,26 | 11,6,6,5,7,7,8,7,10,- | | | |
| 10182021_ Osa_01 | | | | | | 7 | 20 | 12 |
| 10182021_ Osa_02 | | | 7 | 27,27,27 | 6,6,5,7,7,8,7,11,- | | | |
| 10182021_ Osa_03 | | | | | | 7 | 20 | 12 |
| 10182021_ Osa_04 | 4,5,6,3 | 7,8,8,7,7 | 7 | 27,27,27 | 6,6,5,7,7,8,-,-,- | | | |
| 10182021_ Osa_05 | | | | | | 7 | 20 | 12 |
| 10182021_ Osa_06 | 4,5,6,3 | 8,9,8,7,7 | 7 | 27,27,27 | 6,6,5,7,7,8,7,11,- | | | |
| 10182021_ Osa_07 | | | | | | 7 | 20 | 13 |
| 10182021_ Osa_08 | -,5,6,3 | 8,9,8,7,7 | 7 | 25,25,25 | 6,6,5,7,7,8,7,-,- | | | |
| 10182021_ Osa_09 | 4,5,6,3 | 8,-,-,-,- | 7 | 27,27,27 | | | | |
| 10182021_ Osa_10 | | | | | | 7 | 20 | 12 |
| 10182021_ Osa_11 | 4,5,6,3 | 8,9,8,7,7 | 7 | 25,25,25 | 6,6,5,7,7,7,6,11,- | | | |
| **Photo ID** | **AC1 - AC4** | **AE1 - AE5** | **GF** | **N1 - N3** | **PF1 - PF10** | **EG** | **EY** | **CP** |
| 05172022_ Bar_01 | 4,5,6,3 | 8,9,8,7,6 | 7 | 27,27,27 | 6,6,5,7,7,8,7,11,- | 7 | - | 9 |
| 05172022_ Bar_02 | 4,5,6,3 | 8,9,8,7,7 | 7 | 25,25,25 | 6,6,5,7,7,8,7,11,- | 7 | 20 | 9 |
| 05172022_ Bar_03 | 4,5,6,3 | 8,9,8,-,7 | 7 | 25,25,25 | 6,5,5,7,7,8,7,11,- | 7 | 20 | 8 |
| 05172022_ Bar_04 | 4,5,6,3 | 8,9,8,-,- | 7 | 25,25,25 | 6,5,5,7,7,8,7,11,- | | | |
| 05172022_ Bar_05 | 4,5,6,3 | 8,9,8,6,7 | 7 | 26,26,26 | 6,6,5,7,7,8,7,11,- | 7 | 20 | 9 |

**Supplementary Figure S2** Nuclei staining of the trunk epithelium of (**A**) Okinawa, (**B**) Osaka, and (**C**) Barcelona *O. dioica* laboratory strains

**A**

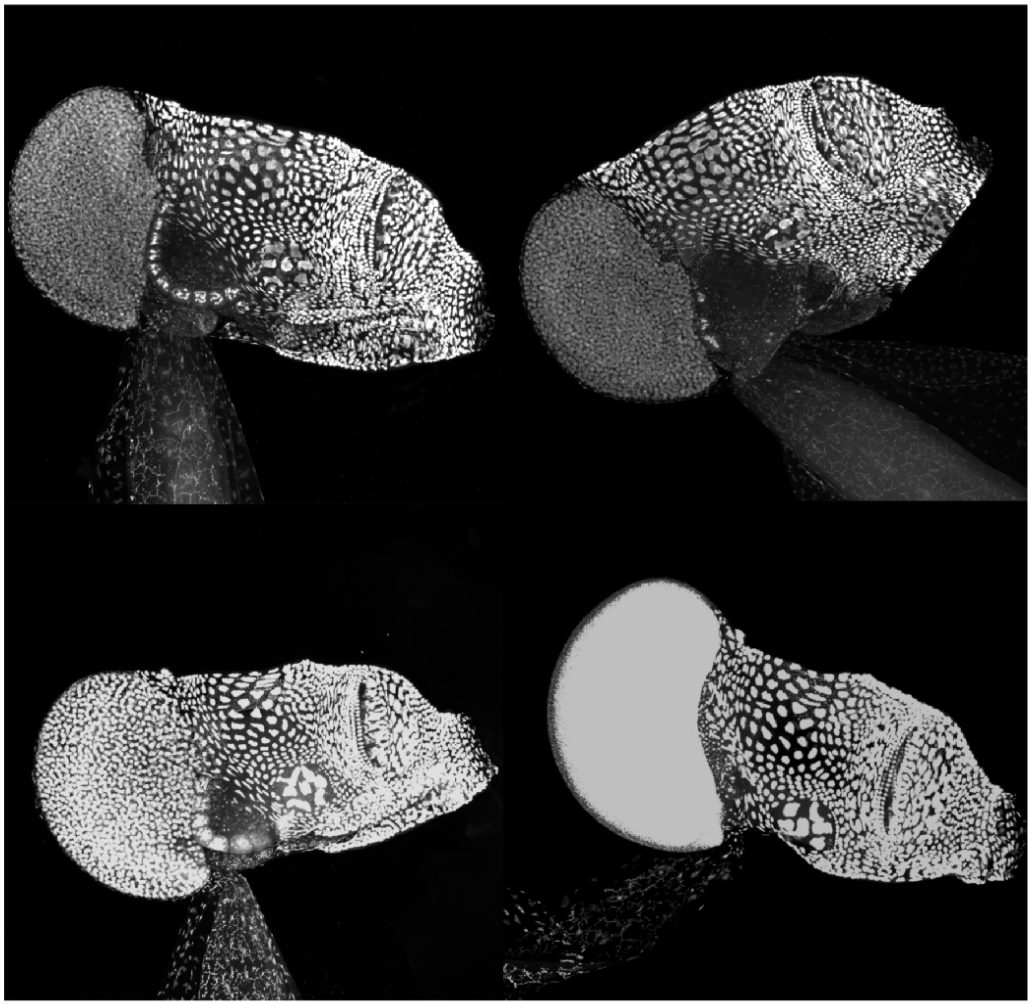

B

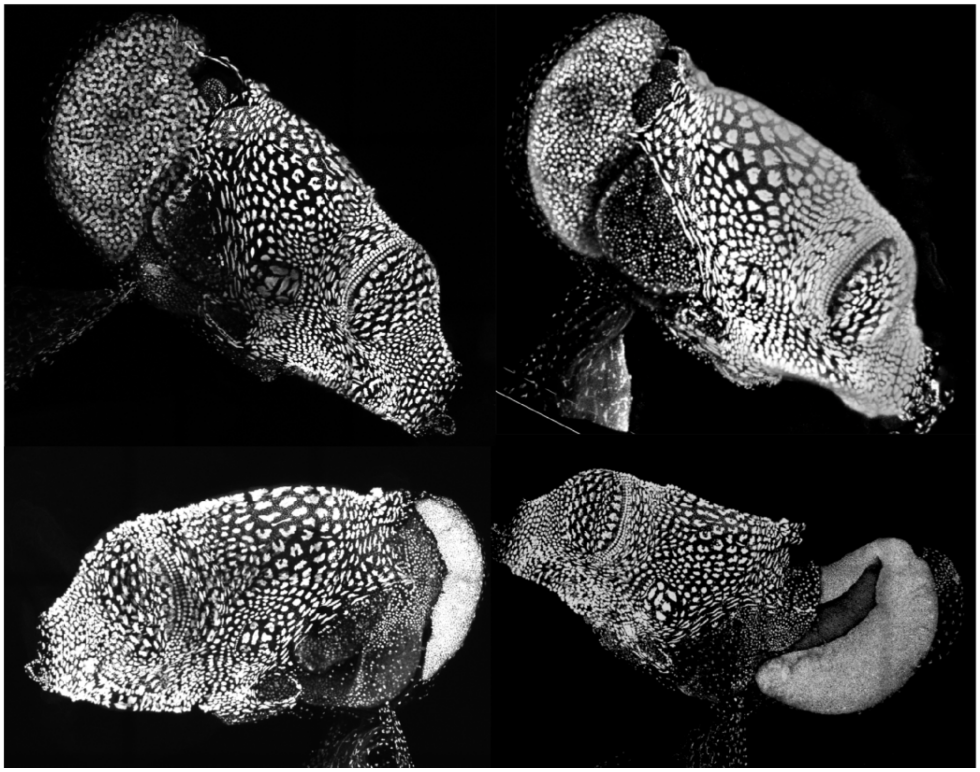

C

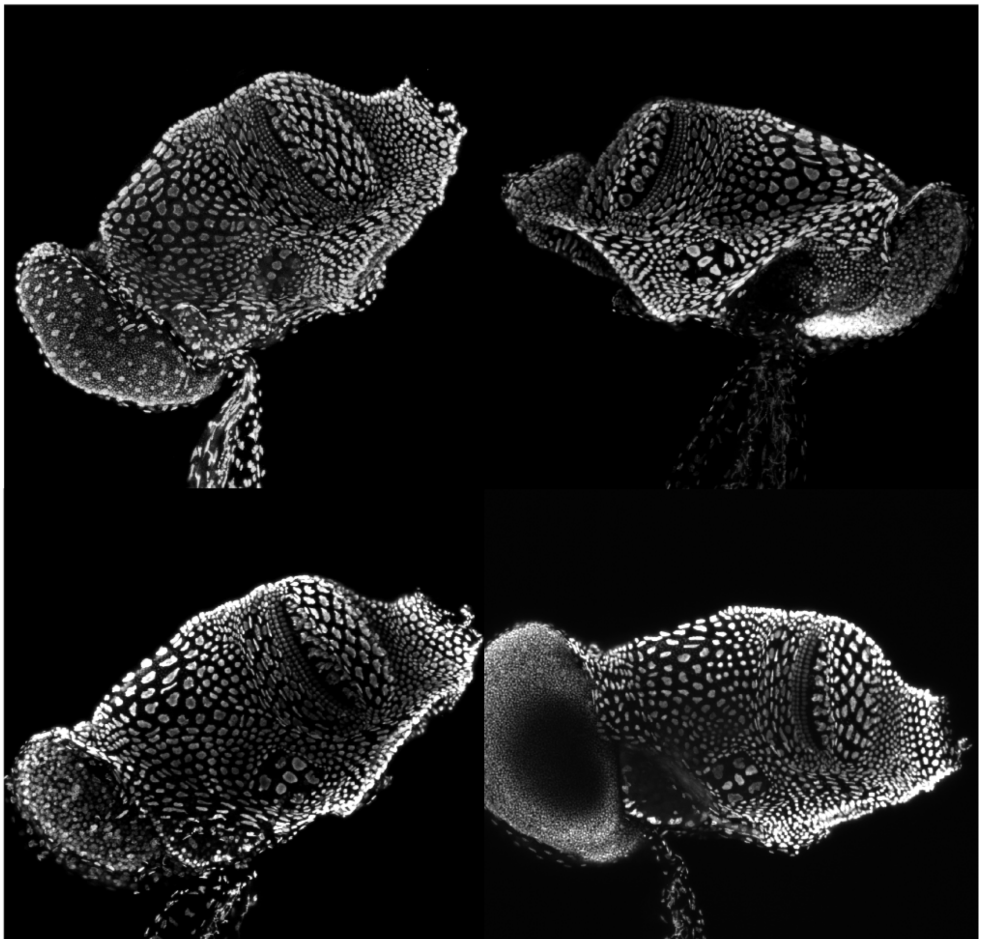

**Supplementary Table S3** Measurements of trunk and tail lengths of Okinawa, Osaka, and Barcelona laboratory strains

| Date | Sample ID | Population | Trunk length (mm) | Tail length (mm) | Ratio (trunk/tail) |
|---|---|---|---|---|---|
| 11182021 | Oki_I23_01 | Kin bay, Okinawa | 0.558 | 2.229 | 0.25 |
| 11182021 | Oki_I23_02 | Kin bay, Okinawa | 0.624 | 2.382 | 0.26 |
| 11182021 | Osa_O10_01 | Sakoshi bay, Hyogo | 0.703 | 2.388 | 0.29 |
| 11182021 | Osa_O10_02 | Sakoshi bay, Hyogo | 0.93 | 2.991 | 0.31 |
| 11222021 | Osa_O10_01 | Sakoshi bay, Hyogo | 0.405 | 1.43 | 0.28 |
| 11222021 | Osa_O10_02 | Sakoshi bay, Hyogo | 0.711 | 2.217 | 0.32 |
| 11222021 | Osa_O10_03 | Sakoshi bay, Hyogo | 0.752 | 2.34 | 0.32 |
| 11222021 | Osa_O10_04 | Sakoshi bay, Hyogo | 0.854 | 2.398 | 0.36 |
| 11222021 | Osa_O10_05 | Sakoshi bay, Hyogo | 0.682 | 2.193 | 0.31 |
| 11222021 | Osa_O10_06 | Sakoshi bay, Hyogo | 0.69 | 2.239 | 0.31 |
| 11222021 | Osa_O10_07 | Sakoshi bay, Hyogo | 0.441 | 1.478 | 0.30 |
| 11222021 | Osa_O10_08 | Sakoshi bay, Hyogo | 0.571 | 1.805 | 0.32 |
| 11222021 | Osa_O10_09 | Sakoshi bay, Hyogo | 0.795 | 2.508 | 0.32 |
| 11222021 | Osa_O10_10 | Sakoshi bay, Hyogo | 0.457 | 1.577 | 0.29 |
| 11252021 | Oki_I23_1 | Kin bay, Okinawa | 0.65 | 2.151 | 0.30 |
| 11252021 | Oki_I23_2 | Kin bay, Okinawa | 0.564 | 2.269 | 0.25 |
| 11252021 | Oki_I23_3 | Kin bay, Okinawa | 0.655 | 2.474 | 0.26 |
| 11252021 | Oki_I23_4 | Kin bay, Okinawa | 0.672 | 2.489 | 0.27 |
| 11252021 | Oki_I23_5 | Kin bay, Okinawa | 0.631 | 2.173 | 0.29 |
| 11252021 | Oki_I23_6 | Kin bay, Okinawa | 0.559 | 2.189 | 0.26 |

| | | | | | |
|---|---|---|---|---|---|
| 11252021 | Oki_I23_7 | Kin bay, Okinawa | 0.627 | 2.13 | 0.29 |
| 11252021 | Oki_I23_8 | Kin bay, Okinawa | 0.397 | 1.66 | 0.24 |
| 11252021 | Oki_I23_9 | Kin bay, Okinawa | 0.504 | 1.941 | 0.26 |
| 11252021 | Oki_I23_10 | Kin bay, Okinawa | 0.386 | 1.49 | 0.26 |
| 11252021 | Oki_I23_11 | Kin bay, Okinawa | 0.561 | 2.226 | 0.25 |
| 11292021 | Osa_O10_01 | Sakoshi bay, Hyogo | 0.485 | 1.777 | 0.27 |
| 11292021 | Osa_O10_02 | Sakoshi bay, Hyogo | 0.854 | 2.583 | 0.33 |
| 11292021 | Osa_O10_03 | Sakoshi bay, Hyogo | 0.781 | 2.671 | 0.29 |
| 11292021 | Osa_O10_04 | Sakoshi bay, Hyogo | 0.586 | 2.149 | 0.27 |
| 11292021 | Osa_O10_05 | Sakoshi bay, Hyogo | 0.541 | 2.046 | 0.26 |
| 11292021 | Osa_O10_06 | Sakoshi bay, Hyogo | 0.604 | 2.027 | 0.30 |
| 11292021 | Osa_O10_07 | Sakoshi bay, Hyogo | 0.869 | 2.695 | 0.32 |
| 11292021 | Osa_O10_08 | Sakoshi bay, Hyogo | 0.553 | 2.084 | 0.27 |
| 11292021 | Osa_O10_09 | Sakoshi bay, Hyogo | 0.54 | 2.072 | 0.26 |
| 11292021 | Osa_O10_10 | Sakoshi bay, Hyogo | 0.53 | 1.965 | 0.27 |
| 12022021 | Oki_I23_01 | Kin bay, Okinawa | 0.501 | 2.148 | 0.23 |
| 12022021 | Oki_I23_02 | Kin bay, Okinawa | 0.723 | 2.358 | 0.31 |
| 12022021 | Oki_I23_03 | Kin bay, Okinawa | 0.645 | 2.086 | 0.31 |
| 12022021 | Oki_I23_04 | Kin bay, Okinawa | 0.801 | 2.612 | 0.31 |
| 12022021 | Oki_I23_05 | Kin bay, Okinawa | 0.662 | 2.523 | 0.26 |
| 12022021 | Oki_I23_06 | Kin bay, Okinawa | 0.505 | 2.039 | 0.25 |
| 12022021 | Oki_I23_07 | Kin bay, Okinawa | 0.474 | 1.535 | 0.31 |
| 12022021 | Oki_I23_08 | Kin bay, Okinawa | 0.547 | 2.186 | 0.25 |

| | | | | | |
|---|---|---|---|---|---|
| 12022021 | Oki_I23_09 | Kin bay, Okinawa | 0.578 | 2.227 | 0.26 |
| 12022021 | Oki_I23_10 | Kin bay, Okinawa | 0.52 | 1.985 | 0.26 |
| 12032021 | Osa_O10_01 | Sakoshi bay, Hyogo | 0.764 | 2.694 | 0.28 |
| 12032021 | Osa_O10_02 | Sakoshi bay, Hyogo | 0.55 | 1.791 | 0.31 |
| 12032021 | Osa_O10_03 | Sakoshi bay, Hyogo | 0.671 | 2.069 | 0.32 |
| 12032021 | Osa_O10_04 | Sakoshi bay, Hyogo | 0.814 | 2.507 | 0.32 |
| 12032021 | Osa_O10_05 | Sakoshi bay, Hyogo | 0.932 | 2.723 | 0.34 |
| 12032021 | Osa_O10_06 | Sakoshi bay, Hyogo | 0.672 | 2.178 | 0.31 |
| 12032021 | Osa_O10_07 | Sakoshi bay, Hyogo | 0.553 | 1.89 | 0.29 |
| 12032021 | Osa_O10_08 | Sakoshi bay, Hyogo | 0.628 | 2.14 | 0.29 |
| 12032021 | Osa_O10_09 | Sakoshi bay, Hyogo | 0.928 | 2.996 | 0.31 |
| 12032021 | Osa_O10_10 | Sakoshi bay, Hyogo | 0.607 | 2.046 | 0.30 |
| 12082021 | Oki_I23_01 | Kin bay, Okinawa | 0.579 | 2.194 | 0.26 |
| 12082021 | Oki_I23_02 | Kin bay, Okinawa | 0.523 | 1.997 | 0.26 |
| 12082021 | Oki_I23_03 | Kin bay, Okinawa | 0.676 | 2.429 | 0.28 |
| 12082021 | Oki_I23_04 | Kin bay, Okinawa | 0.622 | 2.429 | 0.26 |
| 12082021 | Oki_I23_05 | Kin bay, Okinawa | 0.595 | 2.129 | 0.28 |
| 12082021 | Oki_I23_06 | Kin bay, Okinawa | 0.554 | 2.11 | 0.26 |
| 12082021 | Oki_I23_07 | Kin bay, Okinawa | 0.51 | 2.093 | 0.24 |
| 12082021 | Oki_I23_08 | Kin bay, Okinawa | 0.54 | 2.117 | 0.26 |
| 12082021 | Oki_I23_09 | Kin bay, Okinawa | 0.583 | 2.215 | 0.26 |
| 12082021 | Oki_I23_10 | Kin bay, Okinawa | 0.534 | 2.14 | 0.25 |
| 12092021 | Osa_O10_01 | Sakoshi bay, Hyogo | 0.899 | 2.589 | 0.35 |

| | | | | | |
|---|---|---|---|---|---|
| 12092021 | Osa_O10_02 | Sakoshi bay, Hyogo | 0.687 | 2.109 | 0.33 |
| 12092021 | Osa_O10_03 | Sakoshi bay, Hyogo | 0.517 | 1.767 | 0.29 |
| 12092021 | Osa_O10_04 | Sakoshi bay, Hyogo | 0.689 | 2.24 | 0.31 |
| 12092021 | Osa_O10_05 | Sakoshi bay, Hyogo | 1.06 | 2.749 | 0.39 |
| 12092021 | Osa_O10_06 | Sakoshi bay, Hyogo | 0.523 | 1.753 | 0.30 |
| 12092021 | Osa_O10_07 | Sakoshi bay, Hyogo | 0.507 | 1.787 | 0.28 |
| 12092021 | Osa_O10_08 | Sakoshi bay, Hyogo | 0.594 | 1.985 | 0.30 |
| 12092021 | Osa_O10_09 | Sakoshi bay, Hyogo | 0.968 | 2.669 | 0.36 |
| 12092021 | Osa_O10_10 | Sakoshi bay, Hyogo | 0.784 | 2.413 | 0.32 |
| 12102021 | Oki_I23_01 | Kin bay, Okinawa | 0.575 | 2.038 | 0.28 |
| 12102021 | Oki_I23_02 | Kin bay, Okinawa | 0.687 | 2.13 | 0.32 |
| 12102021 | Oki_I23_03 | Kin bay, Okinawa | 0.771 | 2.486 | 0.31 |
| 12102021 | Oki_I23_04 | Kin bay, Okinawa | 0.567 | 2.082 | 0.27 |
| 12102021 | Oki_I23_05 | Kin bay, Okinawa | 0.584 | 1.984 | 0.29 |
| 12102021 | Oki_I23_06 | Kin bay, Okinawa | 0.762 | 2.545 | 0.30 |
| 12102021 | Oki_I23_07 | Kin bay, Okinawa | 0.72 | 2.287 | 0.31 |
| 12102021 | Oki_I23_08 | Kin bay, Okinawa | 0.726 | 2.539 | 0.29 |
| 12102021 | Oki_I23_09 | Kin bay, Okinawa | 0.543 | 1.915 | 0.28 |
| 12102021 | Oki_I23_10 | Kin bay, Okinawa | 0.583 | 2.105 | 0.28 |
| 12132021 | Osa_O10_01 | Sakoshi bay, Hyogo | 0.702 | 2.231 | 0.31 |
| 12132021 | Osa_O10_02 | Sakoshi bay, Hyogo | 0.556 | 1.688 | 0.33 |
| 12132021 | Osa_O10_03 | Sakoshi bay, Hyogo | 0.546 | 1.582 | 0.35 |
| 12132021 | Osa_O10_04 | Sakoshi bay, Hyogo | 0.848 | 2.503 | 0.34 |

| | | | | | |
|---|---|---|---|---|---|
| 12132021 | Osa_O10_05 | Sakoshi bay, Hyogo | 0.935 | 2.603 | 0.36 |
| 12132021 | Osa_O10_06 | Sakoshi bay, Hyogo | 0.694 | 1.997 | 0.35 |
| 12132021 | Osa_O10_07 | Sakoshi bay, Hyogo | 0.86 | 2.694 | 0.32 |
| 12132021 | Osa_O10_08 | Sakoshi bay, Hyogo | 0.863 | 2.653 | 0.33 |
| 12132021 | Osa_O10_09 | Sakoshi bay, Hyogo | 0.853 | 2.605 | 0.33 |
| 12132021 | Osa_O10_10 | Sakoshi bay, Hyogo | 0.741 | 2.241 | 0.33 |
| 12142021 | Oki_I23_01 | Kin bay, Okinawa | 0.411 | 2.041 | 0.20 |
| 12142021 | Oki_I23_02 | Kin bay, Okinawa | 0.471 | 1.856 | 0.25 |
| 12142021 | Oki_I23_03 | Kin bay, Okinawa | 0.387 | 1.567 | 0.25 |
| 12142021 | Oki_I23_04 | Kin bay, Okinawa | 0.584 | 2.218 | 0.26 |
| 12142021 | Oki_I23_05 | Kin bay, Okinawa | 0.499 | 1.943 | 0.26 |
| 12142021 | Oki_I23_06 | Kin bay, Okinawa | 0.558 | 1.96 | 0.28 |
| 12142021 | Oki_I23_07 | Kin bay, Okinawa | 0.441 | 1.591 | 0.28 |
| 12142021 | Oki_I23_08 | Kin bay, Okinawa | 0.55 | 2.148 | 0.26 |
| 12142021 | Oki_I23_09 | Kin bay, Okinawa | 0.483 | 1.637 | 0.30 |
| 12142021 | Oki_I23_10 | Kin bay, Okinawa | 0.419 | 1.671 | 0.25 |
| 05122022 | Bar_01 | Barcelona | 0.75 | 3.09 | 0.243 |
| 05122022 | Bar_02 | Barcelona | 0.65 | 2.71 | 0.240 |
| 05122022 | Bar_03 | Barcelona | 0.67 | 2.52 | 0.266 |
| 05122022 | Bar_04 | Barcelona | 0.68 | 2.34 | 0.291 |
| 05122022 | Bar_05 | Barcelona | 0.681 | 2.53 | 0.269 |
| 05122022 | Bar_06 | Barcelona | 0.782 | 2.88 | 0.271 |
| 05122022 | Bar_07 | Barcelona | 0.702 | 2.59 | 0.271 |

| | | | | | |
|---|---|---|---|---|---|
| 05122022 | Bar_08 | Barcelona | 0.781 | 3.112 | 0.251 |
| 05122022 | Bar_09 | Barcelona | 0.78 | 2.69 | 0.290 |
| 05122022 | Bar_10 | Barcelona | 0.77 | 2.7 | 0.285 |
| 05122022 | Bar_11 | Barcelona | 0.77 | 2.74 | 0.281 |
| 05122022 | Bar_12 | Barcelona | 0.62 | 2.55 | 0.243 |
| 05122022 | Bar_13 | Barcelona | 0.64 | 2.5 | 0.256 |
| 05122022 | Bar_14 | Barcelona | 0.6 | 2.32 | 0.259 |
| 05122022 | Bar_15 | Barcelona | 0.71 | 2.71 | 0.262 |
| 05122022 | Bar_16 | Barcelona | 0.62 | 2.53 | 0.245 |
| 05122022 | Bar_17 | Barcelona | 0.68 | 2.65 | 0.257 |
| 05122022 | Bar_18 | Barcelona | 0.52 | 2.01 | 0.259 |
| 05122022 | Bar_19 | Barcelona | 0.59 | 2.37 | 0.249 |
| 05122022 | Bar_20 | Barcelona | 0.58 | 2.16 | 0.269 |
| 05122022 | Bar_21 | Barcelona | 0.53 | 2.08 | 0.255 |
| 05122022 | Bar_22 | Barcelona | 0.64 | 2.39 | 0.268 |
| 05122022 | Bar_23 | Barcelona | 0.53 | 2.06 | 0.257 |
| 05122022 | Bar_24 | Barcelona | 0.6 | 2.33 | 0.258 |
| 05122022 | Bar_25 | Barcelona | 0.69 | 2.35 | 0.294 |
| 05122022 | Bar_26 | Barcelona | 0.46 | 1.8 | 0.256 |
| 05122022 | Bar_27 | Barcelona | 0.64 | 2.31 | 0.277 |
| 05122022 | Bar_28 | Barcelona | 0.58 | 2.32 | 0.250 |
| 05122022 | Bar_29 | Barcelona | 0.51 | 1.89 | 0.270 |
| 05122022 | Bar_30 | Barcelona | 0.45 | 1.69 | 0.266 |

| | | | | | |
|---|---|---|---|---|---|
| 05122022 | Bar_31 | Barcelona | 0.51 | 1.98 | 0.258 |
| 05122022 | Bar_32 | Barcelona | 0.53 | 2.05 | 0.259 |
| 05122022 | Bar_33 | Barcelona | 0.5 | 2.07 | 0.242 |
| 05122022 | Bar_34 | Barcelona | 0.48 | 1.96 | 0.245 |
| 05122022 | Bar_35 | Barcelona | 0.45 | 1.79 | 0.251 |
| 05122022 | Bar_36 | Barcelona | 0.44 | 1.71 | 0.257 |
| 05122022 | Bar_37 | Barcelona | 0.4 | 1.68 | 0.238 |
| 05122022 | Bar_38 | Barcelona | 0.45 | 1.84 | 0.245 |
| 05122022 | Bar_39 | Barcelona | 0.42 | 1.67 | 0.251 |
| 05122022 | Bar_40 | Barcelona | 0.6 | 1.86 | 0.323 |
| 05122022 | Bar_41 | Barcelona | 0.44 | 1.62 | 0.272 |
| 05122022 | Bar_42 | Barcelona | 0.38 | 1.49 | 0.255 |
| 05122022 | Bar_43 | Barcelona | 0.38 | 1.51 | 0.252 |
| 05122022 | Bar_44 | Barcelona | 0.43 | 1.84 | 0.234 |
| 05122022 | Bar_45 | Barcelona | 0.45 | 1.76 | 0.256 |
| 05122022 | Bar_46 | Barcelona | 0.46 | 1.58 | 0.291 |
| 05122022 | Bar_47 | Barcelona | 0.39 | 1.64 | 0.238 |
| 05122022 | Bar_48 | Barcelona | 0.3 | 1.08 | 0.278 |
| 05122022 | Bar_49 | Barcelona | 0.28 | 1.12 | 0.250 |
| 05122022 | Bar_50 | Barcelona | 0.24 | 0.84 | 0.286 |

**Supplementary Table S4** Egg diameter measurements of Okinawa, Osaka, and Barcelona laboratory strains

| Date | Sample ID | Population | Egg diameter (mm) |
|---|---|---|---|
| 10062021 | I23_1 | Kin bay, Okinawa | 0.081 |
| 10062021 | I23_1 | Kin bay, Okinawa | 0.079 |
| 10062021 | I23_1 | Kin bay, Okinawa | 0.078 |
| 10062021 | I23_1 | Kin bay, Okinawa | 0.081 |
| 10062021 | I23_1 | Kin bay, Okinawa | 0.081 |
| 10062021 | I23_2 | Kin bay, Okinawa | 0.084 |
| 10062021 | I23_2 | Kin bay, Okinawa | 0.085 |
| 10062021 | I23_2 | Kin bay, Okinawa | 0.082 |
| 10062021 | I23_2 | Kin bay, Okinawa | 0.085 |
| 10062021 | I23_2 | Kin bay, Okinawa | 0.082 |
| 10062021 | I23_3 | Kin bay, Okinawa | 0.082 |
| 10062021 | I23_3 | Kin bay, Okinawa | 0.078 |
| 10062021 | I23_3 | Kin bay, Okinawa | 0.081 |
| 10062021 | I23_3 | Kin bay, Okinawa | 0.08 |
| 10062021 | I23_3 | Kin bay, Okinawa | 0.08 |
| 10062021 | O10_1 | Sakoshi bay, Hyogo | 0.097 |
| 10062021 | O10_1 | Sakoshi bay, Hyogo | 0.094 |
| 10062021 | O10_1 | Sakoshi bay, Hyogo | 0.094 |
| 10062021 | O10_1 | Sakoshi bay, Hyogo | 0.096 |
| 10062021 | O10_1 | Sakoshi bay, Hyogo | 0.091 |
| 11052021 | Oki_I23_1 | Kin bay, Okinawa | 0.078 |

| | | | |
|---|---|---|---|
| 11052021 | Oki_I23_1 | Kin bay, Okinawa | 0.079 |
| 11052021 | Oki_I23_1 | Kin bay, Okinawa | 0.081 |
| 11052021 | Oki_I23_1 | Kin bay, Okinawa | 0.077 |
| 11052021 | Oki_I23_1 | Kin bay, Okinawa | 0.08 |
| 11052021 | Oki_I23_1 | Kin bay, Okinawa | 0.081 |
| 11052021 | Oki_I23_02 | Kin bay, Okinawa | 0.084 |
| 11052021 | Oki_I23_02 | Kin bay, Okinawa | 0.086 |
| 11052021 | Oki_I23_02 | Kin bay, Okinawa | 0.086 |
| 11052021 | Oki_I23_02 | Kin bay, Okinawa | 0.081 |
| 11052021 | Oki_I23_02 | Kin bay, Okinawa | 0.084 |
| 11052021 | Osa_O10_1 | Sakoshi bay, Hyogo | 0.094 |
| 11052021 | Osa_O10_1 | Sakoshi bay, Hyogo | 0.095 |
| 11052021 | Osa_O10_1 | Sakoshi bay, Hyogo | 0.098 |
| 11052021 | Osa_O10_1 | Sakoshi bay, Hyogo | 0.097 |
| 11052021 | Osa_O10_1 | Sakoshi bay, Hyogo | 0.098 |
| 11052021 | Osa_O10_2 | Sakoshi bay, Hyogo | 0.098 |
| 11052021 | Osa_O10_2 | Sakoshi bay, Hyogo | 0.097 |
| 11052021 | Osa_O10_2 | Sakoshi bay, Hyogo | 0.101 |
| 11052021 | Osa_O10_2 | Sakoshi bay, Hyogo | 0.097 |
| 11052021 | Osa_O10_2 | Sakoshi bay, Hyogo | 0.098 |
| 11052021 | Osa_O10_3 | Sakoshi bay, Hyogo | 0.101 |
| 11052021 | Osa_O10_3 | Sakoshi bay, Hyogo | 0.1 |
| 11052021 | Osa_O10_3 | Sakoshi bay, Hyogo | 0.1 |

| | | | |
|---|---|---|---|
| 11052021 | Osa_O10_3 | Sakoshi bay, Hyogo | 0.099 |
| 11052021 | Osa_O10_3 | Sakoshi bay, Hyogo | 0.102 |
| 11082021 | Oki_I23_01 | Kin bay, Okinawa | 0.083 |
| 11082021 | Oki_I23_01 | Kin bay, Okinawa | 0.081 |
| 11082021 | Oki_I23_01 | Kin bay, Okinawa | 0.079 |
| 11082021 | Oki_I23_01 | Kin bay, Okinawa | 0.081 |
| 11082021 | Oki_I23_01 | Kin bay, Okinawa | 0.082 |
| 11082021 | Oki_I23_02 | Kin bay, Okinawa | 0.084 |
| 11082021 | Oki_I23_02 | Kin bay, Okinawa | 0.084 |
| 11082021 | Oki_I23_02 | Kin bay, Okinawa | 0.082 |
| 11082021 | Oki_I23_02 | Kin bay, Okinawa | 0.084 |
| 11082021 | Oki_I23_02 | Kin bay, Okinawa | 0.084 |
| 11082021 | Oki_I23_03 | Kin bay, Okinawa | 0.078 |
| 11082021 | Oki_I23_03 | Kin bay, Okinawa | 0.077 |
| 11082021 | Oki_I23_03 | Kin bay, Okinawa | 0.077 |
| 11082021 | Oki_I23_03 | Kin bay, Okinawa | 0.078 |
| 11082021 | Oki_I23_03 | Kin bay, Okinawa | 0.08 |
| 11082021 | Osa_O10_01 | Sakoshi bay, Hyogo | 0.1 |
| 11082021 | Osa_O10_01 | Sakoshi bay, Hyogo | 0.099 |
| 11082021 | Osa_O10_01 | Sakoshi bay, Hyogo | 0.096 |
| 11082021 | Osa_O10_01 | Sakoshi bay, Hyogo | 0.098 |
| 11082021 | Osa_O10_01 | Sakoshi bay, Hyogo | 0.097 |
| 11082021 | Osa_O10_02 | Sakoshi bay, Hyogo | 0.099 |

| | | | |
|---|---|---|---|
| 11082021 | Osa_O10_02 | Sakoshi bay, Hyogo | 0.096 |
| 11082021 | Osa_O10_02 | Sakoshi bay, Hyogo | 0.099 |
| 11082021 | Osa_O10_02 | Sakoshi bay, Hyogo | 0.098 |
| 11082021 | Osa_O10_02 | Sakoshi bay, Hyogo | 0.096 |
| 11082021 | Osa_O10_03 | Sakoshi bay, Hyogo | 0.091 |
| 11082021 | Osa_O10_03 | Sakoshi bay, Hyogo | 0.098 |
| 11082021 | Osa_O10_03 | Sakoshi bay, Hyogo | 0.097 |
| 11082021 | Osa_O10_03 | Sakoshi bay, Hyogo | 0.098 |
| 11082021 | Osa_O10_03 | Sakoshi bay, Hyogo | 0.095 |
| 11092021 | Oki_I23_01 | Kin bay, Okinawa | 0.075 |
| 11092021 | Oki_I23_01 | Kin bay, Okinawa | 0.074 |
| 11092021 | Oki_I23_01 | Kin bay, Okinawa | 0.075 |
| 11092021 | Oki_I23_01 | Kin bay, Okinawa | 0.072 |
| 11092021 | Oki_I23_01 | Kin bay, Okinawa | 0.074 |
| 11092021 | Oki_I23_02 | Kin bay, Okinawa | 0.079 |
| 11092021 | Oki_I23_02 | Kin bay, Okinawa | 0.078 |
| 11092021 | Oki_I23_02 | Kin bay, Okinawa | 0.078 |
| 11092021 | Oki_I23_02 | Kin bay, Okinawa | 0.08 |
| 11092021 | Oki_I23_02 | Kin bay, Okinawa | 0.077 |
| 11092021 | Oki_I23_03 | Kin bay, Okinawa | 0.081 |
| 11092021 | Oki_I23_03 | Kin bay, Okinawa | 0.083 |
| 11092021 | Oki_I23_03 | Kin bay, Okinawa | 0.079 |
| 11092021 | Oki_I23_03 | Kin bay, Okinawa | 0.081 |

| | | | |
|---|---|---|---|
| 11092021 | Oki_I23_03 | Kin bay, Okinawa | 0.081 |
| 11092021 | Osa_O10_01 | Sakoshi bay, Hyogo | 0.097 |
| 11092021 | Osa_O10_01 | Sakoshi bay, Hyogo | 0.095 |
| 11092021 | Osa_O10_01 | Sakoshi bay, Hyogo | 0.096 |
| 11092021 | Osa_O10_01 | Sakoshi bay, Hyogo | 0.1 |
| 11092021 | Osa_O10_01 | Sakoshi bay, Hyogo | 0.098 |
| 11092021 | Osa_O10_02 | Sakoshi bay, Hyogo | 0.095 |
| 11092021 | Osa_O10_02 | Sakoshi bay, Hyogo | 0.095 |
| 11092021 | Osa_O10_02 | Sakoshi bay, Hyogo | 0.093 |
| 11092021 | Osa_O10_02 | Sakoshi bay, Hyogo | 0.094 |
| 11092021 | Osa_O10_02 | Sakoshi bay, Hyogo | 0.095 |
| 11092021 | Osa_O10_03 | Sakoshi bay, Hyogo | 0.091 |
| 11092021 | Osa_O10_03 | Sakoshi bay, Hyogo | 0.092 |
| 11092021 | Osa_O10_03 | Sakoshi bay, Hyogo | 0.09 |
| 11092021 | Osa_O10_03 | Sakoshi bay, Hyogo | 0.09 |
| 11092021 | Osa_O10_03 | Sakoshi bay, Hyogo | 0.091 |
| 11102021 | Oki_I23_01 | Kin bay, Okinawa | 0.078 |
| 11102021 | Oki_I23_01 | Kin bay, Okinawa | 0.076 |
| 11102021 | Oki_I23_01 | Kin bay, Okinawa | 0.078 |
| 11102021 | Oki_I23_01 | Kin bay, Okinawa | 0.078 |
| 11102021 | Oki_I23_01 | Kin bay, Okinawa | 0.078 |
| 11102021 | Oki_I23_02 | Kin bay, Okinawa | 0.078 |
| 11102021 | Oki_I23_02 | Kin bay, Okinawa | 0.076 |

| | | | |
|---|---|---|---|
| 11102021 | Oki_I23_02 | Kin bay, Okinawa | 0.078 |
| 11102021 | Oki_I23_02 | Kin bay, Okinawa | 0.078 |
| 11102021 | Oki_I23_02 | Kin bay, Okinawa | 0.078 |
| 11102021 | Oki_I23_03 | Kin bay, Okinawa | 0.081 |
| 11102021 | Oki_I23_03 | Kin bay, Okinawa | 0.082 |
| 11102021 | Oki_I23_03 | Kin bay, Okinawa | 0.082 |
| 11102021 | Oki_I23_03 | Kin bay, Okinawa | 0.083 |
| 11102021 | Oki_I23_03 | Kin bay, Okinawa | 0.082 |
| 11102021 | Osa_O10_01 | Sakoshi bay, Hyogo | 0.091 |
| 11102021 | Osa_O10_01 | Sakoshi bay, Hyogo | 0.09 |
| 11102021 | Osa_O10_01 | Sakoshi bay, Hyogo | 0.088 |
| 11102021 | Osa_O10_01 | Sakoshi bay, Hyogo | 0.09 |
| 11102021 | Osa_O10_01 | Sakoshi bay, Hyogo | 0.089 |
| 11102021 | Osa_O10_02 | Sakoshi bay, Hyogo | 0.094 |
| 11102021 | Osa_O10_02 | Sakoshi bay, Hyogo | 0.097 |
| 11102021 | Osa_O10_02 | Sakoshi bay, Hyogo | 0.097 |
| 11102021 | Osa_O10_02 | Sakoshi bay, Hyogo | 0.096 |
| 11102021 | Osa_O10_02 | Sakoshi bay, Hyogo | 0.094 |
| 11112021 | Oki_I23_01 | Kin bay, Okinawa | 0.077 |
| 11112021 | Oki_I23_01 | Kin bay, Okinawa | 0.079 |
| 11112021 | Oki_I23_01 | Kin bay, Okinawa | 0.079 |
| 11112021 | Oki_I23_01 | Kin bay, Okinawa | 0.077 |
| 11112021 | Oki_I23_01 | Kin bay, Okinawa | 0.078 |

| | | | |
|---|---|---|---|
| 11112021 | Oki_I23_02 | Kin bay, Okinawa | 0.078 |
| 11112021 | Oki_I23_02 | Kin bay, Okinawa | 0.078 |
| 11112021 | Oki_I23_02 | Kin bay, Okinawa | 0.077 |
| 11112021 | Oki_I23_02 | Kin bay, Okinawa | 0.077 |
| 11112021 | Oki_I23_02 | Kin bay, Okinawa | 0.077 |
| 11112021 | Oki_I23_03 | Kin bay, Okinawa | 0.082 |
| 11112021 | Oki_I23_03 | Kin bay, Okinawa | 0.08 |
| 11112021 | Oki_I23_03 | Kin bay, Okinawa | 0.08 |
| 11112021 | Oki_I23_03 | Kin bay, Okinawa | 0.081 |
| 11112021 | Oki_I23_03 | Kin bay, Okinawa | 0.082 |
| 11112021 | Osa_O10_01 | Sakoshi bay, Hyogo | 0.099 |
| 11112021 | Osa_O10_01 | Sakoshi bay, Hyogo | 0.098 |
| 11112021 | Osa_O10_01 | Sakoshi bay, Hyogo | 0.1 |
| 11112021 | Osa_O10_01 | Sakoshi bay, Hyogo | 0.098 |
| 11112021 | Osa_O10_01 | Sakoshi bay, Hyogo | 0.099 |
| 11112021 | Osa_O10_02 | Sakoshi bay, Hyogo | 0.103 |
| 11112021 | Osa_O10_02 | Sakoshi bay, Hyogo | 0.098 |
| 11112021 | Osa_O10_02 | Sakoshi bay, Hyogo | 0.098 |
| 11112021 | Osa_O10_02 | Sakoshi bay, Hyogo | 0.098 |
| 11112021 | Osa_O10_02 | Sakoshi bay, Hyogo | 0.1 |
| 11112021 | Osa_O10_03 | Sakoshi bay, Hyogo | 0.096 |
| 11112021 | Osa_O10_03 | Sakoshi bay, Hyogo | 0.096 |
| 11112021 | Osa_O10_03 | Sakoshi bay, Hyogo | 0.098 |

| | | | |
|---|---|---|---|
| 11112021 | Osa_O10_03 | Sakoshi bay, Hyogo | 0.098 |
| 11112021 | Osa_O10_03 | Sakoshi bay, Hyogo | 0.1 |
| 11112021 | Osa_O10_04 | Sakoshi bay, Hyogo | 0.099 |
| 11112021 | Osa_O10_04 | Sakoshi bay, Hyogo | 0.097 |
| 11112021 | Osa_O10_04 | Sakoshi bay, Hyogo | 0.1 |
| 11112021 | Osa_O10_04 | Sakoshi bay, Hyogo | 0.1 |
| 11112021 | Osa_O10_04 | Sakoshi bay, Hyogo | 0.098 |
| 11122021 | Oki_I23_01 | Kin bay, Okinawa | 0.079 |
| 11122021 | Oki_I23_01 | Kin bay, Okinawa | 0.077 |
| 11122021 | Oki_I23_01 | Kin bay, Okinawa | 0.082 |
| 11122021 | Oki_I23_01 | Kin bay, Okinawa | 0.081 |
| 11122021 | Oki_I23_01 | Kin bay, Okinawa | 0.082 |
| 11122021 | Oki_I23_02 | Kin bay, Okinawa | 0.076 |
| 11122021 | Oki_I23_02 | Kin bay, Okinawa | 0.074 |
| 11122021 | Oki_I23_02 | Kin bay, Okinawa | 0.075 |
| 11122021 | Oki_I23_02 | Kin bay, Okinawa | 0.077 |
| 11122021 | Oki_I23_02 | Kin bay, Okinawa | 0.075 |
| 11122021 | Oki_I23_03 | Kin bay, Okinawa | 0.08 |
| 11122021 | Oki_I23_03 | Kin bay, Okinawa | 0.081 |
| 11122021 | Oki_I23_03 | Kin bay, Okinawa | 0.081 |
| 11122021 | Oki_I23_03 | Kin bay, Okinawa | 0.08 |
| 11122021 | Oki_I23_03 | Kin bay, Okinawa | 0.079 |
| 11122021 | Osa_O10_01 | Sakoshi bay, Hyogo | 0.09 |

| | | | |
|---|---|---|---|
| 11122021 | Osa_O10_01 | Sakoshi bay, Hyogo | 0.091 |
| 11122021 | Osa_O10_01 | Sakoshi bay, Hyogo | 0.09 |
| 11122021 | Osa_O10_01 | Sakoshi bay, Hyogo | 0.089 |
| 11122021 | Osa_O10_01 | Sakoshi bay, Hyogo | 0.09 |
| 11122021 | Osa_O10_02 | Sakoshi bay, Hyogo | 0.093 |
| 11122021 | Osa_O10_02 | Sakoshi bay, Hyogo | 0.095 |
| 11122021 | Osa_O10_02 | Sakoshi bay, Hyogo | 0.095 |
| 11122021 | Osa_O10_02 | Sakoshi bay, Hyogo | 0.092 |
| 11122021 | Osa_O10_02 | Sakoshi bay, Hyogo | 0.094 |
| 11122021 | Osa_O10_03 | Sakoshi bay, Hyogo | 0.09 |
| 11122021 | Osa_O10_03 | Sakoshi bay, Hyogo | 0.091 |
| 11122021 | Osa_O10_03 | Sakoshi bay, Hyogo | 0.09 |
| 11122021 | Osa_O10_03 | Sakoshi bay, Hyogo | 0.09 |
| 11122021 | Osa_O10_03 | Sakoshi bay, Hyogo | 0.091 |
| 11172021 | Osa_O10_01 | Sakoshi bay, Hyogo | 0.096 |
| 11172021 | Osa_O10_01 | Sakoshi bay, Hyogo | 0.095 |
| 11172021 | Osa_O10_01 | Sakoshi bay, Hyogo | 0.096 |
| 11172021 | Osa_O10_01 | Sakoshi bay, Hyogo | 0.096 |
| 11172021 | Osa_O10_01 | Sakoshi bay, Hyogo | 0.097 |
| 11172021 | Osa_O10_02 | Sakoshi bay, Hyogo | 0.097 |
| 11172021 | Osa_O10_02 | Sakoshi bay, Hyogo | 0.096 |
| 11172021 | Osa_O10_02 | Sakoshi bay, Hyogo | 0.099 |
| 11172021 | Osa_O10_02 | Sakoshi bay, Hyogo | 0.098 |

| | | | |
|---|---|---|---|
| 11172021 | Osa_O10_02 | Sakoshi bay, Hyogo | 0.098 |
| 05122022 | Bar_F1 | Barcelona | 0.079 |
| 05122022 | Bar_F1 | Barcelona | 0.083 |
| 05122022 | Bar_F1 | Barcelona | 0.082 |
| 05122022 | Bar_F1 | Barcelona | 0.082 |
| 05122022 | Bar_F1 | Barcelona | 0.082 |
| 05122022 | Bar_F2 | Barcelona | 0.091 |
| 05122022 | Bar_F2 | Barcelona | 0.093 |
| 05122022 | Bar_F2 | Barcelona | 0.093 |
| 05122022 | Bar_F2 | Barcelona | 0.095 |
| 05122022 | Bar_F2 | Barcelona | 0.101 |
| 05122022 | Bar_F3 | Barcelona | 0.102 |
| 05122022 | Bar_F3 | Barcelona | 0.094 |
| 05122022 | Bar_F3 | Barcelona | 0.101 |
| 05122022 | Bar_F3 | Barcelona | 0.104 |
| 05122022 | Bar_F3 | Barcelona | 0.108 |
| 05122022 | Bar_F4 | Barcelona | 0.087 |
| 05122022 | Bar_F4 | Barcelona | 0.083 |
| 05122022 | Bar_F4 | Barcelona | 0.088 |
| 05122022 | Bar_F4 | Barcelona | 0.088 |
| 05122022 | Bar_F4 | Barcelona | 0.092 |
| 05122022 | Bar_F5 | Barcelona | 0.12 |
| 05122022 | Bar_F5 | Barcelona | 0.096 |

| | | | |
|---|---|---|---|
| 05122022 | Bar_F5 | Barcelona | 0.092 |
| 05122022 | Bar_F5 | Barcelona | 0.119 |
| 05122022 | Bar_F5 | Barcelona | 0.118 |
| 05122022 | Bar_F6 | Barcelona | 0.098 |
| 05122022 | Bar_F6 | Barcelona | 0.098 |
| 05122022 | Bar_F6 | Barcelona | 0.092 |
| 05122022 | Bar_F6 | Barcelona | 0.094 |
| 05122022 | Bar_F6 | Barcelona | 0.096 |
| 05122022 | Bar_F7 | Barcelona | 0.082 |
| 05122022 | Bar_F7 | Barcelona | 0.086 |
| 05122022 | Bar_F7 | Barcelona | 0.087 |
| 05122022 | Bar_F7 | Barcelona | 0.085 |
| 05122022 | Bar_F7 | Barcelona | 0.084 |
| 05122022 | Bar_F8 | Barcelona | 0.079 |
| 05122022 | Bar_F8 | Barcelona | 0.087 |
| 05122022 | Bar_F8 | Barcelona | 0.088 |
| 05122022 | Bar_F8 | Barcelona | 0.086 |
| 05122022 | Bar_F8 | Barcelona | 0.074 |
| 05122022 | Bar_F9 | Barcelona | 0.103 |
| 05122022 | Bar_F9 | Barcelona | 0.095 |
| 05122022 | Bar_F9 | Barcelona | 0.108 |
| 05122022 | Bar_F9 | Barcelona | 0.103 |
| 05122022 | Bar_F9 | Barcelona | 0.106 |

| | | | |
|---|---|---|---|
| 05122022 | Bar_F10 | Barcelona | 0.099 |
| 05122022 | Bar_F10 | Barcelona | 0.102 |
| 05122022 | Bar_F10 | Barcelona | 0.092 |
| 05122022 | Bar_F10 | Barcelona | 0.104 |
| 05122022 | Bar_F10 | Barcelona | 0.099 |
| 05122022 | Bar_F11 | Barcelona | 0.121 |
| 05122022 | Bar_F11 | Barcelona | 0.111 |
| 05122022 | Bar_F11 | Barcelona | 0.119 |
| 05122022 | Bar_F11 | Barcelona | 0.115 |
| 05122022 | Bar_F11 | Barcelona | 0.115 |
| 05122022 | Bar_F12 | Barcelona | 0.101 |
| 05122022 | Bar_F12 | Barcelona | 0.1 |
| 05122022 | Bar_F12 | Barcelona | 0.099 |
| 05122022 | Bar_F12 | Barcelona | 0.1 |
| 05122022 | Bar_F12 | Barcelona | 0.095 |
| 05122022 | Bar_F13 | Barcelona | 0.097 |
| 05122022 | Bar_F13 | Barcelona | 0.099 |
| 05122022 | Bar_F13 | Barcelona | 0.096 |
| 05122022 | Bar_F13 | Barcelona | 0.093 |
| 05122022 | Bar_F13 | Barcelona | 0.095 |
| 05122022 | Bar_F14 | Barcelona | 0.094 |
| 05122022 | Bar_F14 | Barcelona | 0.098 |
| 05122022 | Bar_F14 | Barcelona | 0.093 |

| | | | |
|---|---|---|---|
| 05122022 | Bar_F14 | Barcelona | 0.097 |
| 05122022 | Bar_F14 | Barcelona | 0.094 |
| 05122022 | Bar_F15 | Barcelona | 0.095 |
| 05122022 | Bar_F15 | Barcelona | 0.095 |
| 05122022 | Bar_F15 | Barcelona | 0.086 |
| 05122022 | Bar_F15 | Barcelona | 0.102 |
| 05122022 | Bar_F15 | Barcelona | 0.093 |
| 05122022 | Bar_F16 | Barcelona | 0.102 |
| 05122022 | Bar_F16 | Barcelona | 0.102 |
| 05122022 | Bar_F16 | Barcelona | 0.104 |
| 05122022 | Bar_F16 | Barcelona | 0.096 |
| 05122022 | Bar_F16 | Barcelona | 0.104 |
| 05122022 | Bar_F17 | Barcelona | 0.094 |
| 05122022 | Bar_F17 | Barcelona | 0.089 |
| 05122022 | Bar_F17 | Barcelona | 0.091 |
| 05122022 | Bar_F17 | Barcelona | 0.093 |
| 05122022 | Bar_F17 | Barcelona | 0.091 |
| 05122022 | Bar_F18 | Barcelona | 0.103 |
| 05122022 | Bar_F18 | Barcelona | 0.1 |
| 05122022 | Bar_F18 | Barcelona | 0.105 |
| 05122022 | Bar_F18 | Barcelona | 0.086 |
| 05122022 | Bar_F18 | Barcelona | 0.088 |
| 05122022 | Bar_F19 | Barcelona | 0.096 |

| | | | |
|---|---|---|---|
| 05122022 | Bar_F19 | Barcelona | 0.102 |
| 05122022 | Bar_F19 | Barcelona | 0.098 |
| 05122022 | Bar_F19 | Barcelona | 0.095 |
| 05122022 | Bar_F19 | Barcelona | 0.093 |
| 05122022 | Bar_F20 | Barcelona | 0.091 |
| 05122022 | Bar_F20 | Barcelona | 0.097 |
| 05122022 | Bar_F20 | Barcelona | 0.103 |
| 05122022 | Bar_F20 | Barcelona | 0.092 |
| 05122022 | Bar_F20 | Barcelona | 0.091 |
| 05122022 | Bar_F21 | Barcelona | 0.099 |
| 05122022 | Bar_F21 | Barcelona | 0.096 |
| 05122022 | Bar_F21 | Barcelona | 0.095 |
| 05122022 | Bar_F21 | Barcelona | 0.094 |
| 05122022 | Bar_F21 | Barcelona | 0.097 |

**Supplementary Table 5** Crossing experiments for Okinawa and Osaka laboratory strains

| Date | Female | Male | Time fertilized | # of eggs fertilized |
|---|---|---|---|---|
| 04202018 | Osaka_2 | Okinawa_1 | 14:05 | 0/32 |
| 04202018 | Okinawa_3 | Osaka_1 | 14:05 | 0/34 |
| 04202018 | Osaka_1 | Osaka_2 | 14:05 | 34/35 |
| 04202018 | Okinawa_2 | Osaka_3 | 14:05 | 0/34 |
| 04202018 | Osaka_3 | Okinawa_2 | 14:05 | 0/35 |
| 04202018 | Okinawa_1 | Okinawa_3 | 14:05 | 25/36 |
| 04252018 | Okinawa_3 | Okinawa_2 | 13:30 | 31/34 |
| 04252018 | Osaka_2 | Osaka_3 | 13:30 | 36/36 |
| 04252018 | Osaka_3 | Okinawa_3 | 13:30 | 0/34 |
| 04252018 | Okinawa_2 | Osaka_1 | 13:30 | 0/35 |
| 04252018 | Osaka_1 | Okinawa_1 | 13:30 | 0/37 |
| 04252018 | Okinawa_1 | Osaka_2 | 13:30 | 0/34 |
| 05142018 | Okinawa_3 | Okinawa_2 | 13:18 | 32/32 |
| 05142018 | Osaka_2 | Osaka_3 | 13:18 | 37/39 |
| 05142018 | Osaka_3 | Okinawa_3 | 13:18 | 0/39 |
| 05142018 | Okinawa_2 | Osaka_1 | 13:18 | 0/30 |
| 05142018 | Osaka_1 | Okinawa_1 | 13:18 | 0/44 |
| 05142018 | Okinawa_1 | Osaka_2 | 13:18 | 0/34 |
| 05252018 | Okinawa_3 | Okinawa_2 | 12:45 | 32/35 |
| 05252018 | Osaka_2 | Osaka_3 | 12:45 | 33/33 |
| 05252018 | Osaka_3 | Okinawa_3 | 12:45 | 0/35 |

| | | | | |
|---|---|---|---|---|
| 05252018 | Okinawa_2 | Osaka_1 | 12:45 | 0/31 |
| 05252018 | Osaka_1 | Okinawa_1 | 12:45 | 0/35 |
| 05252018 | Okinawa_1 | Osaka_2 | 12:45 | 0/41 |
| 05302018 | Osaka_1 | Okinawa_2 | 13:24 | 0/42 |
| 05302018 | Okinawa_1 | Osaka_3 | 13:24 | 0/36 |
| 05302018 | Okinawa_2 | Okinawa_3 | 13:24 | 38/42 |
| 05302018 | Okinawa_3 | Osaka_1 | 13:24 | 0/46 |
| 05302018 | Osaka_3 | Okinawa_1 | 13:24 | 0/45 |
| 05302018 | Osaka_2 | Osaka_2 | 13:24 | 54/54 |
